# Mitochondrial antiviral-signalling protein is a client of the BAG6 protein quality control complex

**DOI:** 10.1101/2021.12.02.470791

**Authors:** Peristera Roboti, Craig Lawless, Stephen High

## Abstract

The heterotrimeric BAG6 complex coordinates the direct handover of newly synthesised tail-anchored (TA) membrane proteins from an SGTA-bound preloading complex to the endoplasmic reticulum (ER) delivery component TRC40. In contrast, defective precursors, including aberrant TA proteins, form a stable complex with this cytosolic protein quality control factor, enabling such clients to be either productively re-routed or selectively degraded. We identify the mitochondrial TA protein MAVS (mitochondrial antiviral-signalling protein) as an endogenous client of both SGTA and the BAG6 complex. Our data suggest that the BAG6 complex binds to a cytosolic pool of MAVS before its misinsertion into the ER membrane, from where it can subsequently be removed via ATP13A1-mediated dislocation. This BAG6- associated fraction of MAVS is dynamic and responds to the activation of an innate immune response, suggesting that BAG6 may modulate the pool of MAVS that is available for coordinating the cellular response to viral infection.

**SUMMARY STATEMENT:** Mitochondrial antiviral-signalling (MAVS) protein is a favoured client of the cytosolic BAG6 complex. We discuss how this dynamic interaction may modulate MAVS biogenesis at signalling membranes.

## INTRODUCTION

The endoplasmic reticulum (ER) is a major site of membrane protein synthesis, typically involving the targeting of nascent polypeptide chains to the ER by virtue of hydrophobic targeting signals such as transmembrane domains (TMDs) or cleavable N-terminal signal peptides (Cross et al., 2009). In eukaryotes, many such targeting signals are bound by the cytosolic signal recognition particle (SRP) as they emerge from the ribosomal exit tunnel; thereby enabling their co-translational delivery to the ER for subsequent membrane insertion (Cross et al., 2009; O’Keefe et al., 2021a; O’Keefe et al., 2021b). In addition to such well characterised co-translational pathways (O’Keefe et al., 2021a), several post-translational routes for the delivery of completed membrane proteins to the ER have been described (Farkas and Bohnsack, 2021; Hegde and Keenan, 2021). Furthermore, recent studies have revealed that, in addition to supplying the compartments of the secretory pathway and plasma membrane with newly synthesised membrane proteins, the ER may also act as a staging post for membrane proteins *en route* to mitochondria. Hence, membrane proteins that initially mislocalise to the ER may be redirected to mitochondria via a mechanism termed ER-SURF (Koch et al., 2021).

Tail-anchored (TA) membrane proteins are characterised by the presence of a single TMD at their extreme C-terminus (Kutay et al., 1993), which necessitates their post-translational delivery to the appropriate subcellular organelle (Farkas and Bohnsack, 2021; Johnson et al., 2013). Whilst most TA proteins are inserted into the ER membrane, a smaller group is delivered to the mitochondrial outer membrane (MOM) (Costello et al., 2017). Although multiple redundant pathways have been identified, many mammalian TA proteins are delivered to the ER via the TMD recognition complex (TRC) targeting pathway (Casson et al., 2017; Farkas and Bohnsack, 2021; Hegde and Keenan, 2021). Here, SGTA captures TA clients shortly after their TMDs emerge from the ribosome to form a so-called pretargeting complex (Farkas and Bohnsack, 2021; Leznicki and High, 2020). Subsequently, the binding of SGTA to the heterotrimeric Bag6-Ubl4A-TRC35 (hereafter BAG6) complex enables TA proteins to be handed off to the ER targeting factor TRC40 (Farkas and Bohnsack, 2021). Once bound to TRC40, TA proteins are delivered to the ER by binding to the WRB-CAML complex, which acts as both a membrane receptor and insertase (McDowell et al., 2020). In comparison to our detailed understanding of TA protein biogenesis at the ER (Casson et al., 2017; Farkas and Bohnsack, 2021; Hegde and Keenan, 2021), the components and mechanisms responsible for TA protein insertion into other organellar membranes are less well-defined (Farkas and Bohnsack, 2021). One suggestion is that the membrane lipid composition may be an important factor in the selective insertion of TA proteins into the MOM (Brambillasca et al., 2005; Krumpe et al., 2012), whilst in yeast, Pex19p has been implicated in the integration of certain TA proteins at both the MOM and peroxisomes (Cichocki et al., 2018; Mayerhofer, 2016).

SGTA functions as a homodimeric cochaperone comprised of three functional domains (see also Fig. 1A). The N-terminal region acts as a homodimerisation module that, once assembled, enables the SGTA dimer to bind either the Ubl4A or Bag6 subunits of the BAG6 complex via their respective N-terminal ubiquitin-like (UBL) domains (Roberts et al., 2015). The central tetratricopeptide repeat (TPR) domain of SGTA interacts with cytosolic heat-shock proteins and the proteasomal subunit Rpn13 (Roberts et al., 2015). The C-terminal methionine-rich domain of SGTA binds hydrophobic TMDs (Liou and Wang, 2005). Its flexibility and dimeric nature are suggested to facilitate the shielding of hydrophobic TA regions and enable conformational changes that mediate downstream interactions (Lin et al., 2021; Martinez-Lumbreras et al., 2018).

**Fig. 1.**
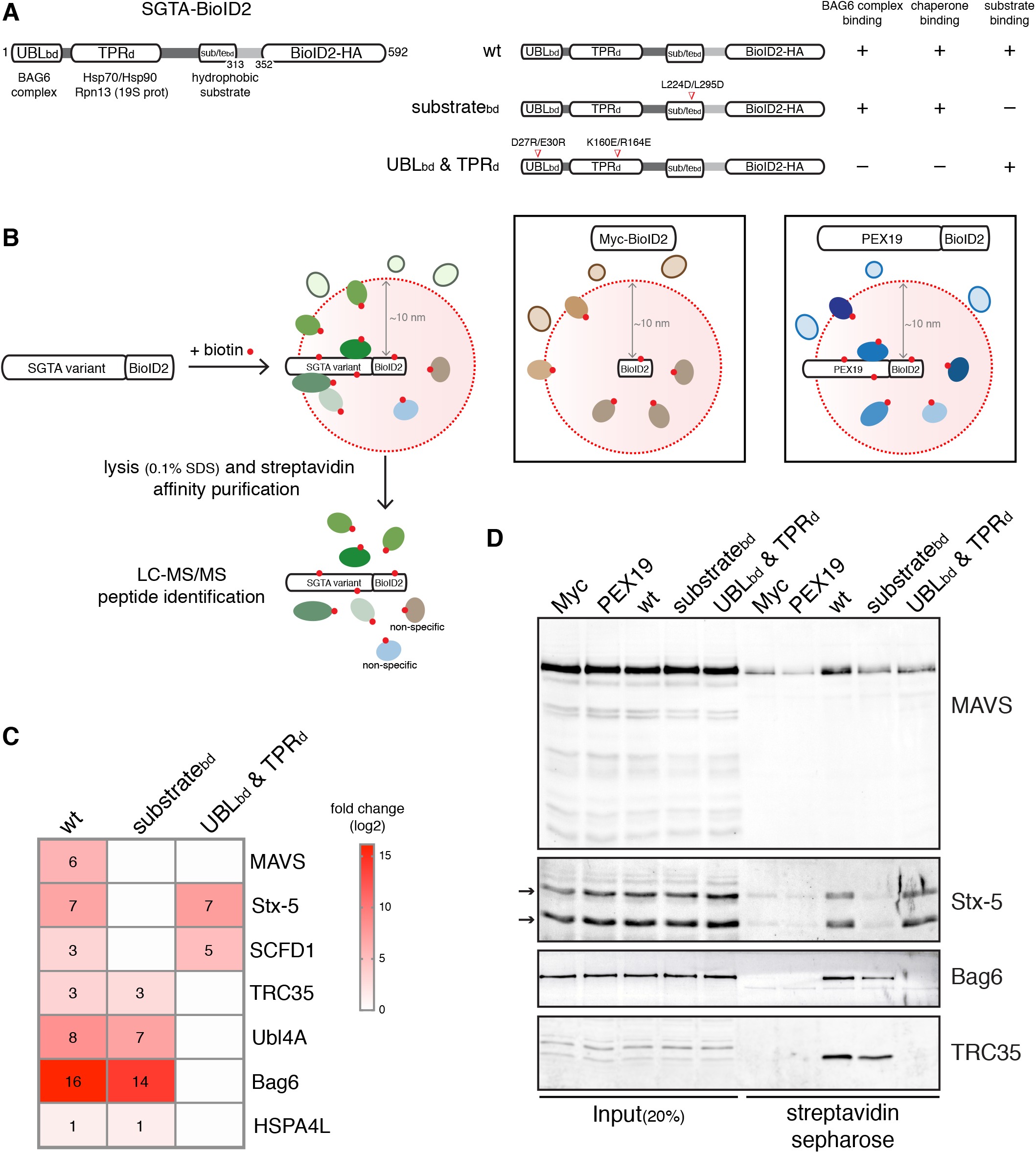
MAVS is a high-confidence proximal interactor of SGTA. (A) Left: Schematic of human SGTA-BioID2-HA displaying its protein-protein interaction modules. UBLbd, N-terminal domain that binds to the ubiquitin-like (UBL) domains of Ubl4A and Bag6; TPRd, central tetratricopeptide repeat (TPR) domain that interacts with heat-shock proteins; substratebd, C-terminal domain that contains the hydrophobic substrate-binding site. Right: Schematics of the SGTA-BioID2 mutants and respective disrupted interactions. Mutated amino acids are indicated. (B) Scheme for the BioID2- mediated proximity labelling and identification of SGTA substrates and cofactors using SGTA KO cells (see Fig. S1) transiently expressing wild-type (wt) SGTA-BioID2 or mutant variants shown in (A). Cells expressing Myc-BioID2 or PEX19-BioID2 serve as two independent controls to exclude non-specific interactors. (C) Heat map representing log2-transformed fold changes in the protein intensities of significant (BFDR<0.05) wild-type (wt)/mutant SGTA-specific preys relative to both the Myc- BioID2 and PEX19-BioID2 controls. Individual rounded values are depicted in the heat map. A non-significant prey is shown as a white box (three biological replicates, see Table S1 for list of all proteins detected). (D) Validation of selected SGTA- associated candidates from (C) by immunoblotting. SGTA KO cells expressing the indicated BioID2-tagged baits were treated with biotin for 8 h and lysed with RIPA buffer. The resulting extracts were subjected to affinity purification with streptavidin beads and the bound material eluted using a biotin-containing buffer. The input and eluted material were analysed by immunoblotting for the indicated endogenous proteins. Stx-5 can be observed as two bands (hereafter indicated by arrows) corresponding to two isoforms, a 42 kDa-ER and a 35 kDa-Golgi isoform that result from an alternative initiation of translation (Coy-Vergara et al., 2019).

The high-affinity binding of SGTA to Ubl4A subunit of the BAG6 complex facilitates the prompt and privileged transfer of TA protein clients from SGTA directly to TRC40 (Mock et al., 2015; Shao et al., 2017). In contrast, the lower affinity interaction between SGTA and the Bag6 subunit (Krysztofinska et al., 2016) enables mislocalised or defective hydrophobic precursor proteins including aberrant TA proteins (hereafter termed mislocalised proteins (MLPs)) to be passed on from SGTA to the poorly characterised central region of the Bag6 subunit (Leznicki et al., 2013). The capacity of the BAG6 complex to recruit specialised E3 ligases facilitates the subsequent ubiquitination and proteasomal degradation of such MLP clients (Hessa et al., 2011; Rodrigo-Brenni et al., 2014; Shao et al., 2017; Suzuki and Kawahara, 2016; Wunderley et al., 2014; Yamamoto et al., 2017). Interestingly, SGTA can antagonise this BAG6-depedent polyubiquitination, perhaps increasing the window of opportunity for TA protein clients to access a productive outcome (Casson et al., 2016; Leznicki and High, 2012).

Although cells or tissues that lack a functional TRC pathway show substrate-specific defects in the ER targeting and membrane insertion of essential TA proteins, they remain viable (Casson et al., 2017; Jonikas et al., 2009; Norlin et al., 2016; Pfaff et al., 2016; Rivera-Monroy et al., 2016; Schuldiner et al., 2008; Vogl et al., 2016), consistent with the presence of one or more alternative pathways for TA protein biogenesis at the ER (Casson et al., 2017; Hassdenteufel et al., 2017). Likewise, at the ER membrane, in addition to the WRB/CAML-dependent integration of TA proteins delivered via TRC40 (McDowell et al., 2020), the ER membrane protein complex (EMC) acts as an ER membrane insertase for TA proteins with moderately hydrophobic TMDs that cannot efficiently exploit the TRC40 pathway (Guna et al., 2018).

Here, we use BioID2-based proximity labelling to identify the mitochondrial antiviral-signalling protein (MAVS) as an endogenous TA client of SGTA. MAVS has been detected at the MOM, peroxisomes and mitochondria-associated ER membrane subdomains (MAMs) and is implicated in innate immune signalling at each of these locations (Thoresen et al., 2021; Vazquez and Horner, 2015). MAVS is also prone to being “mislocalised” to the ER membrane, from where it can be removed by the P5A-type ATPase ATP13A1 (McKenna et al., 2020). Like other TA proteins, we find that MAVS can be handed off from SGTA to the BAG6 quality control complex, which is stably associated with a pool of cytosolic MAVS. To further explore the origin of this BAG6-bound pool of MAVS, we manipulated its ER-mislocalised form by knocking down its likely membrane insertase, the EMC, and dislocase, the ATP13A1. Our resulting data suggest a model where cytosolic MAVS binds the BAG6 complex before misinsertion into the ER membrane. BAG6-bound MAVS responds to the activation of an innate immune response and we speculate that BAG6 may modulate MAVS activity, perhaps by contributing to the recently identified ER-SURF pathway and/or supplying MAVS to MAMs.

## RESULTS

### BioID2 screening for proximal SGTA interactors

To identify clients and cofactors of SGTA, we used the promiscuous biotin ligase BioID2 to label its neighbouring proteins in cultured mammalian cells (Kim et al., 2016), reasoning this approach would be well suited to identifying relevant weak and/or transient interactions (Shao et al., 2017; Wunderley et al., 2014; Xu et al., 2012). We generated SGTA baits that carried BioID2-HA (hereafter BioID2) at the C-terminus, a position close enough to the substrate-binding domain to allow labelling (Liou and Wang, 2005; Martinez-Lumbreras et al., 2018; Wang et al., 2010). Three forms of SGTA were used; wild-type SGTA, a substrate-binding domain mutant (substrate_bd_mt), exhibiting significantly reduced affinity for hydrophobic clients (Lin et al., 2021) and a combined UBL-binding and TPR domain mutant ((UBL_bd_ & TPR_d_)mt), defective in its interactions with the BAG6 complex, molecular chaperones and the proteasome, but retaining its ability to bind clients (Leznicki et al., 2015; Walczak et al., 2014; Xu et al., 2012) (Fig. 1A).

By comparing the profile of biotinylated proteins obtained using these three SGTA variants, we hoped to distinguish candidate substrates/clients from cofactors. In order to control for the specificity of these SGTA proximal proteomes, we also used cells expressing either Myc-tagged BioID2 or BioID2-HA fused to PEX19, a cytosolic chaperone implicated in the biogenesis of lipid droplet proteins at the ER (Schrul and Kopito, 2016) and TA proteins at peroxisomes and mitochondria (Cichocki et al., 2018; Mayerhofer, 2016) (Fig. 1B). These various BioID2 fusions were expressed in SGTA knockout (KO) HepG2 cells (Fig. S1A,B), thereby removing any competition with the endogenous protein. Although the TA proteins that SGTA hands over to TRC40 most likely include several SNARE proteins that are involved in vesicular trafficking (Coy-Vergara et al., 2019), global protein secretion appeared unaffected in the SGTA KO cell line (Fig. S1C).

In preliminary experiments, BioID2-tagged baits were transiently expressed in SGTA KO cells that were treated with exogenous biotin for 8 h to confirm labelling efficiency. In contrast to the smaller Myc-BioID2 that was expressed throughout the cell, the SGTA-BioID2 variants and PEX19-BioID2 fusion all showed a diffuse cytosolic signal by immunofluorescence microscopy, consistent with the location of biotinylated proteins that were labelled using a streptavidin-conjugated fluorescent probe (Fig. S2A). Likewise, comparable levels of self-biotinylation were seen with each of the BioID2-tagged baits (Fig. S2B) and we proceeded to analyse their respective interactomes. As proof of principle, we next examined the ability of SGTA- BioID2 to biotinylate known interacting partners by immunoblotting (Fig. S2C-E). As expected, the Bag6 protein and Hsp90 were present in the streptavidin-bound material recovered from cells expressing wild-type SGTA-BioID2 and substrate_bd_mt, but were absent in the pull-down material from cells expressing (UBL_bd_ & TPR_d_)mt (Fig. S2C-E). The known TRC40 client syntaxin-5 (Stx-5) (Coy-Vergara et al., 2019; Norlin et al., 2016) was efficiently recovered from the lysate of cells expressing wild-type SGTA-BioID2, but much reduced from that of substrate_bd_mt-expressing cells (Fig. S2C,E). We therefore concluded that we can selectively biotinylate known SGTA partners and discriminate between cofactors and substrates based on their labelling with different SGTA-BioID2 variants.

To further characterise the proximal environment of SGTA, we performed large-scale BioID2 experiments in SGTA KO HepG2 cells, identified high-confidence interactors using the significance analysis of interactome (SAINT) algorithm (Teo et al., 2014) (Fig. 1C; Fig. S3; complete list in Table S1) and validated selected candidates by immunoblotting (Fig. 1D). Wild-type and substrate_bd_mt baits confirmed the proximity of SGTA to the three subunits of the BAG6 complex i.e. Bag6, Ubl4A and TRC35 (see (Casson et al., 2016)) and the Hsp70 chaperone, HSPA4L (see Table S3 in (Pourhaghighi et al., 2020)) (Fig. 1C,D). In contrast, the TRC40 TA protein client Stx-5 (Coy-Vergara et al., 2019) and its conserved partner SCFD1 (Sec1 family domain-containing protein 1) were selectively labelled by wild-type and (UBL_bd_ & TPR_d_)mt baits (Fig. 1C,D). These results validated the data from our small-scale study, identified Stx-5 as a bona fide SGTA client and confirmed that our approach can differentiate between cofactors and substrates.

Our BioID2 analysis suggested a selective association between wild-type SGTA and the mitochondrial TA protein MAVS (see Fig. 1C; Fig. S3; Table S1). Furthermore, biotinylated MAVS was efficiently recovered in the streptavidin pull-down material from cells expressing wild-type SGTA, but was reduced in pull-downs using lysates of cells expressing both SGTA variants (Fig. 1D). The mitochondrial localisation of MAVS, together with its role in innate immunity (Thoresen et al., 2021), prompted us to study its interaction with SGTA in more detail.

### MAVS is a substrate of SGTA

To confirm the findings from our BioID2 labelling, we assessed the association between endogenous SGTA and MAVS by co-immunoprecipitation using cytosol prepared from control KO HepG2 cells (Fig. 2A-C). This fraction was enriched for known cytosolic proteins including SGTA, Bag6 and tubulin, but largely depleted of integral membrane protein markers for the ER (Calnexin) and mitochondria (TOM20), which were recovered in the pellet (Fig. 2A). Interestingly, ∼40% of full-length MAVS was recovered in the cytosolic fraction (Fig. 2A), reminiscent of previous reports that a fraction of the Golgi-resident TA protein Stx-5 was recovered in the cytosol (Coy-Vergara et al., 2019). Furthermore, both of these TA proteins were co-immunoprecipitated with SGTA from this cytosolic fraction (Fig. 2B), confirming the interactions that were suggested by our BioID2 analysis (Fig. 1).

**Fig. 2.**
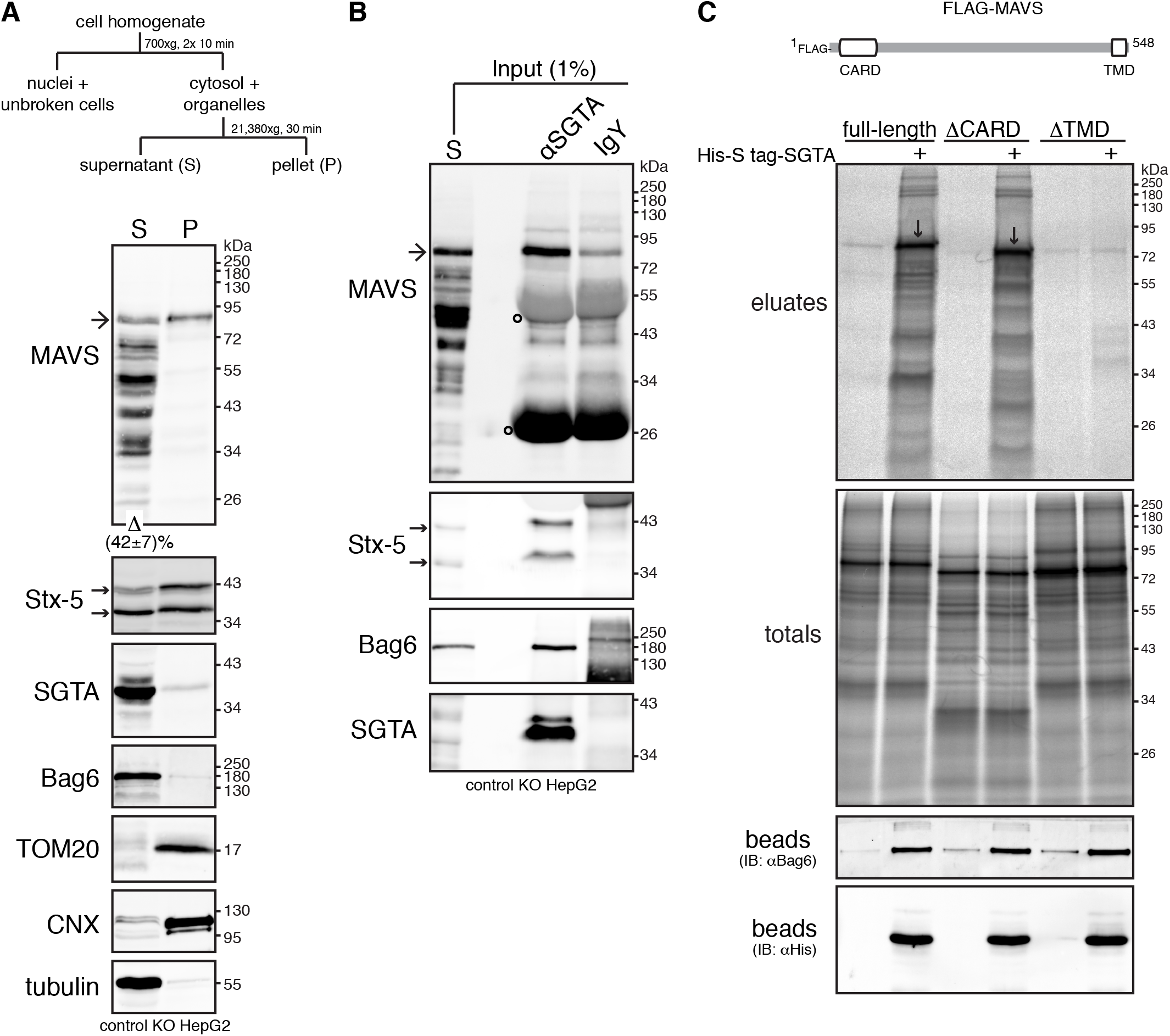
SGTA interacts with MAVS. (A) A cytosolic pool of endogenous MAVS can be observed at steady-state. Top: Schematic of subcellular fractionation protocol used to separate the cell homogenate into crude cytosolic supernatant (S) and membrane-associated pellet (P) fractions. Bottom: Detergent-free extracts from control KO cells (see Fig. S1) were fractionated as shown above. Equivalent amounts of each fraction were analysed by immunoblotting for MAVS and various compartmental markers. Bag6, SGTA and tubulin (cytosolic markers), TOM20 (mitochondrial outer membrane marker) and Calnexin (CNX, ER membrane marker) serve as fractionation controls. Note that the MAVS-specific antibody, raised against amino acids 1-135 of human MAVS, detected the ∼80 kDa full-length MAVS (hereafter marked by an arrow) and multiple shorter variants that most likely represent C-terminally degraded products or processed forms of the full-length protein (see also (Seth et al., 2005)). Quantification of the levels of full-length MAVS recovered in the cytosolic fraction is indicated below the MAVS blot. Value represents mean ± s.e.m. from three independent experiments. (B) MAVS co-immunoprecipitates with SGTA. The supernatant (S) fraction from (A) was subjected to immunoprecipitations with equal amounts of chicken anti-SGTA antibody or chicken IgY antibody (control for non-specific binding). Input and immunoprecipitates were analysed by immunoblotting for the indicated endogenous proteins. Bag6 served as positive control for SGTA binding. Hereafter, open circles on MAVS blots indicate signals derived from denatured antibody heavy and light chains. (C) *In vitro* translated MAVS interacts with recombinant SGTA via its transmembrane domain (TMD). Top: Schematic of FLAG-MAVS displaying its N-terminal caspase activation and recruitment domain (CARD) and C-terminal TMD. Bottom: FLAG-MAVS full-length, ΔCARD or ΔTMD truncated variants were translated *in vitro* in the absence or presence (+) of 2 μM His-S-tag-SGTA. A 10% sample of the total translation products was subjected to denaturing immunoprecipitations with anti-FLAG antibody (totals), while the rest was incubated with HisPur Cobalt resin and bound proteins were eluted using imidazole (eluates). Totals and eluates were resolved by SDS-PAGE and results visualised by phosphorimaging. Downward arrows indicate full-length and ΔCARD FLAG-MAVS selectively bound by His-S-tag-SGTA. His- S-tag-SGTA and its binding partners within rabbit reticulocyte lysate were released from the resin by incubating the beads with SDS sample buffer (beads) and samples were analysed by immunoblotting (IB). The anti-His and anti-Bag6 immunoblots indicate uniform binding of Bag6 binding-competent His-S-tag-SGTA to beads.

Previous *in vitro* studies have shown that SGTA binds to exposed hydrophobic TMDs (Liou and Wang, 2005), including those of TA proteins (Itakura et al., 2016; Leznicki et al., 2010), and we therefore used a pull-down assay (Leznicki and High, 2020) to investigate the interaction of MAVS with SGTA. Following *in vitro* translation of FLAG-tagged MAVS variants in the presence or absence of recombinant His-S-tag- SGTA, SGTA-bound clients were captured using cobalt resin. Radiolabelled wild-type FLAG-MAVS and a version lacking its characteristic N-terminal caspase activation and recruitment domain (ΔCARD) were both efficiently recovered with SGTA (Fig. 2C, eluates). In contrast, removal of its hydrophobic TMD (ΔTMD) prevented MAVS from forming a stable interaction with SGTA (Fig. 2C, eluates), consistent with previous studies of SGTA clients (Leznicki et al., 2010; Leznicki and High, 2020; Wang et al., 2010). Taken together, these results confirm that MAVS is an authentic endogenous binding partner of SGTA and suggest that its primary mode of interaction is TMD dependent.

### MAVS exhibits a robust interaction with the BAG6 complex that is enhanced by SGTA

Previous studies identified MAVS as a potential interacting partner of BAG6 (Antonicka et al., 2020; Li et al., 2011), which typically acts downstream of SGTA (Farkas and Bohnsack, 2021; Hegde and Keenan, 2021). We therefore extended our co-immunoprecipitation study and found that MAVS is also an endogenous client of the BAG6 protein quality control complex (Benarroch et al., 2019) (Fig. 3A; Fig. S4A,B). In contrast to MAVS, we found no evidence that a second endogenous TA protein of the MOM, OMP25, is a potential SGTA client (Table S1) and does co-precipitate with the BAG6 complex (Fig. S4B). Hence, whilst OMP25 was previously shown to interact with SGTA and the BAG6 complex *in vitro* when its target MOM was absent (Itakura et al., 2016), our cell-based assay suggests that alternative TMD-binding factors deal with OMP25 molecules that fail to become membrane-inserted. Furthermore, unlike MAVS, we find no evidence for a substantial pool of OMP25 in the cytosol of HepG2 cells (Fig. S4B). Nevertheless, we explored the possibility that any uninserted OMP25 may bind to the Ubiquilins, which are reported to act as TMD-binding chaperones that prevent OMP25 aggregation prior to its membrane insertion or degradation (Itakura et al., 2016). However, we were unable to confirm any interaction between endogenous OMP25 and Ubiquilin-2 by co-immunoprecipitation (Fig. S5A), consistent with our fractionation studies which suggest that the majority of OMP25 is membrane inserted at steady state (Fig. S4B). Likewise, we find no evidence that MAVS forms a stable complex with the Ubiqulins under identical conditions to those used to recover the BAG6 complex bound to both MAVS and Stx-5 (Fig. S5B). Hence, although MAVS and OMP25 have similarly hydrophobic TMDs and basic residues at their C-termini (Fig. S4C) that are characteristic of TA proteins destined for the MOM (Costello et al., 2017; Rao et al., 2016), our data suggest that MAVS is a favoured client of the BAG6 complex.

**Fig. 3.**
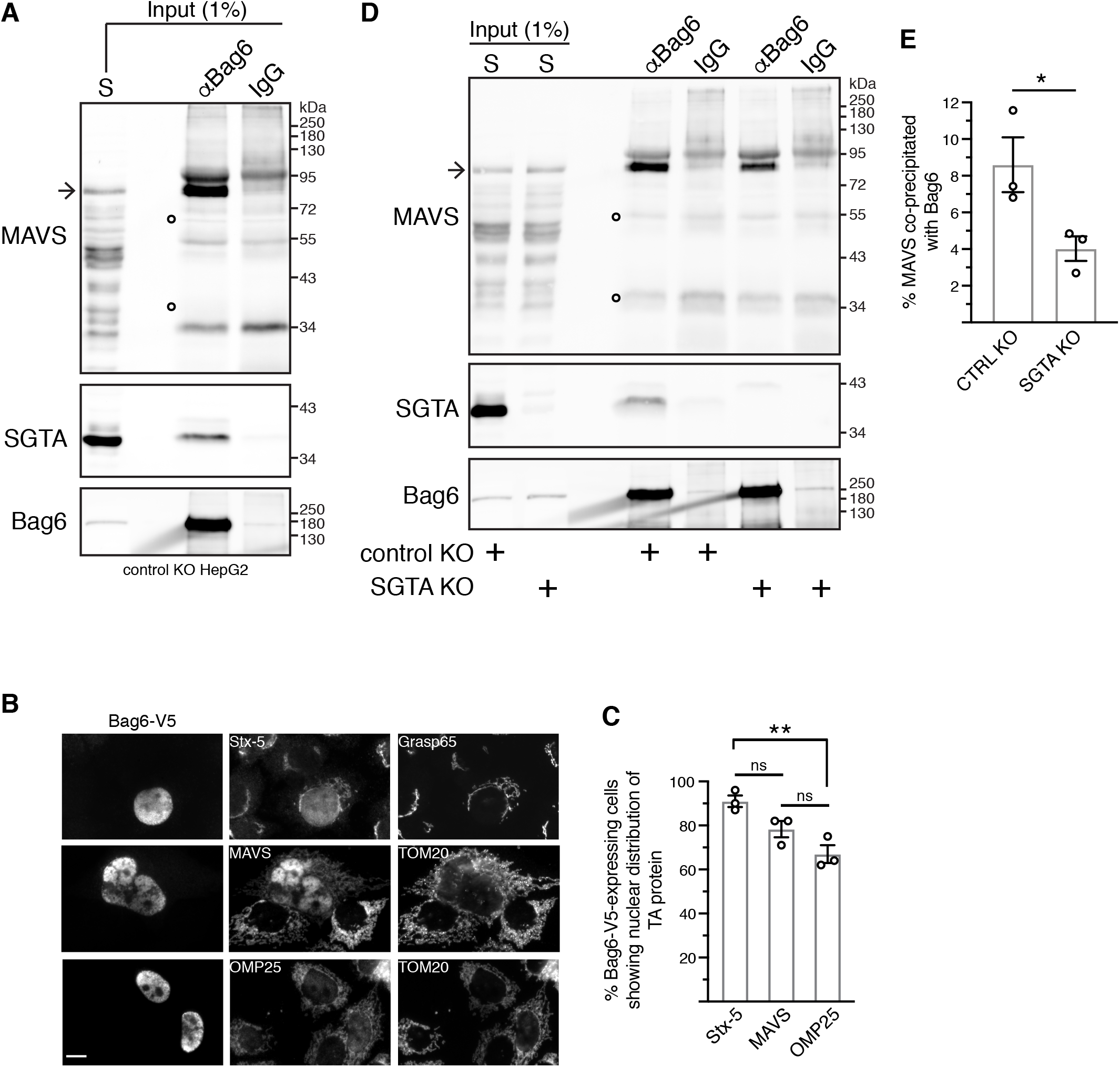
Bag6 interacts with MAVS. (A) MAVS co-immunoprecipitates with Bag6. Control KO cells were fractionated as shown in Fig. 2A and the supernatant (S) fraction was subjected to immunoprecipitations with equal amounts of rabbit anti- Bag6 antibody or rabbit IgG antibody (control for non-specific binding). Input and immunoprecipitates were analysed by immunoblotting for the indicated endogenous proteins. SGTA served as positive control for Bag6 binding. (B) Nuclear redistribution of endogenous TA proteins in Bag6-V5-expressing cells. Representative wide-field fluorescence images of control KO cells transiently transfected with Bag6-V5, fixed and immunostained for exogenous Bag6-V5, the indicated endogenous TA protein and either GRASP65 (*cis*-Golgi marker) or TOM20 (outer mitochondrial membrane marker). Scale bar, 10 μm. (C) Mean ± s.e.m. of number of cells displaying nuclear localisation of the indicated endogenous TA protein. More than 100 cells per condition were analysed from three independent experiments as in (B). **P < 0.01; ns, not significant (ordinary one-way ANOVA with Dunnett’s multiple comparison tests). (D) SGTA facilitates Bag6-MAVS interaction. Control KO and SGTA KO cells were fractionated as shown in Fig. 2A and the supernatant (S) fractions were subjected to immunoprecipitations with rabbit anti-Bag6 antibody or rabbit control IgG antibody. Inputs and immunoprecipitates were analysed by immunoblotting for the indicated endogenous proteins. SGTA served as positive control for Bag6 binding. (E) Mean ± s.e.m. of MAVS levels that co-immunoprecipitate with Bag6 in control KO and SGTA KO cells for three independent experiments as in (D). *P < 0.05 (unpaired two-tailed t test).

To further explore the interaction between BAG6 and TA proteins in a cellular context, we exploited the ability of the Bag6 protein to re-localise hydrophobic clients to the nucleus when exogenously expressed (Leznicki et al., 2010; Payapilly and High, 2014; Wunderley et al., 2014). Hence, whilst ∼90% of cells expressing exogenous Bag6-V5 showed detectable levels of Stx-5 immunofluorescence staining in the nucleus and ∼80% of such cells showed nuclear MAVS staining, significantly fewer cells (<70%) showed evidence of OMP25 re-localisation (Fig. 3B,C). The data from this cell-based assay support a model where the two TA proteins Stx-5 and MAVS are favoured clients of the BAG6 complex.

Our next step was to evaluate whether SGTA played any role in the binding of MAVS to the BAG6 complex, consistent with its previously established role in TA protein biogenesis (Farkas and Bohnsack, 2021). When the amounts of BAG6-bound MAVS recovered from control KO and SGTA KO cells were compared, we observed a ∼50% reduction following SGTA knockout (Fig. 3D,E). These results support previous models, which propose that SGTA acts upstream of the BAG6 complex to which it hands off hydrophobic substrates *en route* to either the ER membrane or regulated proteasomal degradation (Casson et al., 2016; Farkas and Bohnsack, 2021; Shao et al., 2017). However, these data also show that the loss of SGTA does not preclude MAVS binding to BAG6, implying there are alternative pathways for their association (see Discussion). Collectively, these findings suggest that the BAG6 complex interacts with the mitochondrial TA protein MAVS via a mechanism that can be facilitated by SGTA, most likely acting as an upstream delivery factor.

### ATP13A1 deficiency does not affect the BAG6-MAVS interaction

ATP13A1 is an ER-resident P5A-ATPase that has been implicated in the extraction of mitochondrial TA proteins including MAVS and OMP25 that can mislocalise to the ER membrane (McKenna et al., 2020). Given the ability of Bag6 and SGTA to bind membrane protein substrates that are dislocated into the cytosol via the pathways responsible for ERAD (Benarroch et al., 2019), we postulated that BAG6 may bind MAVS after its ATP13A1-mediated dislocation from the ER membrane.

To test this hypothesis, we compared the cytosolic population of MAVS recovered with BAG6 in control and ATP13A1_-_depleted cells, reasoning that an inhibition of ATP13A1_-_mediated dislocation from the ER membrane would impair any downstream interaction of MAVS with the BAG6 complex (Fig. 4A). In ATP13A1- knockdown cells (∼70% reduction; Fig. 4Bi,ii), levels of both total cellular MAVS and its cytoplasmic pool showed a modest decrease (Fig. 4B), consistent with a partially redundant role of ATP13A1 in MAVS dislocation from the ER (McKenna et al., 2020). However, depletion of ATP13A1 had no significant effect on the amount of MAVS that co-immunoprecipitated with the BAG6 complex, which remained stable at ∼20% of the available cytosolic pool of MAVS (Fig. 4C,D). These findings indicate that ATP13A1 deficiency does not compromise the BAG6-bound cohort of MAVS, arguing against the hypothesis that MAVS is handed over to the BAG6 complex via an ATP13A1-dependent mechanism.

**Fig. 4.**
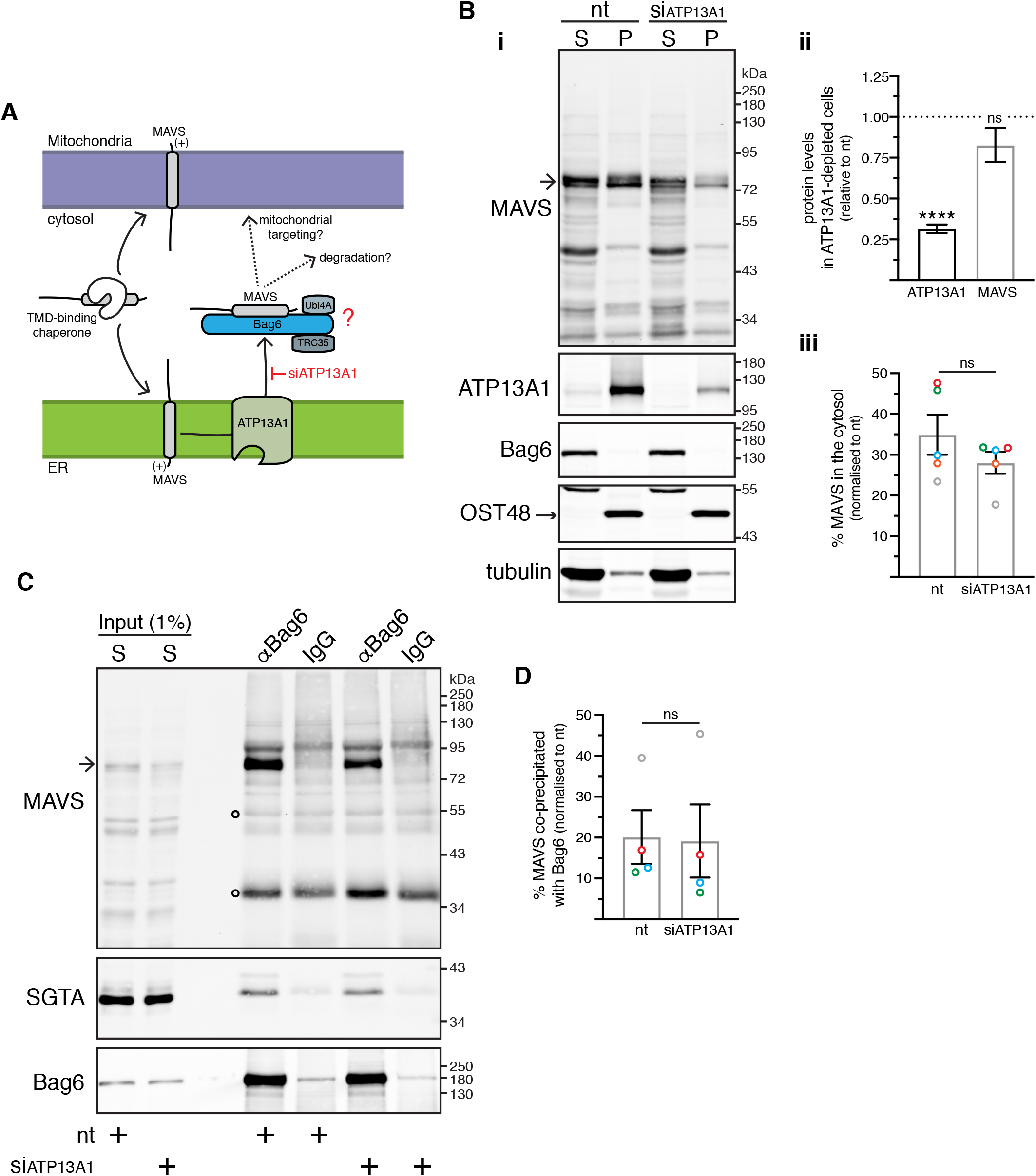
ATP13A1 depletion has no visible effect on Bag6-MAVS interaction. (A) Proposed model: Bag6 recruits MAVS after its ATP13A1-mediated extraction from the ER membrane. Depleting ATP13A1 will decrease Bag6-MAVS interaction, as MAVS cannot be dislocated from the ER membrane. (B) ATP13A1 depletion does not grossly alter the levels of MAVS in the crude cytosolic supernatant fraction. (i) Control KO cells transfected with non-targeting (nt) or ATP13A1-targeting siRNAs were fractionated as shown in Fig. 2A. Equivalent amounts of supernatant (S) and pellet (P) fractions were analysed by immunoblotting for the indicated endogenous proteins. (ii) ATP13A1 and MAVS levels in siATP13A1-treated cells relative to nt siRNA-treated cells, where protein levels were set to 1. Shown are means ± s.e.m. for five biological replicates as shown in (Bi). ****P < 0.0001; ns, not significant (two- tailed one sample t test). (iii) Mean ± s.e.m. of the supernatant/total ratio of MAVS levels in siATP13A1-treated cells normalised to nt siRNA-treated cells for five independent experiments as in (Bi). Same colour data points correspond to a single biological replicate; ns, not significant (paired two-tailed t test). (C) ATP13A1 depletion does not affect Bag6-MAVS interaction. Supernatant (S) fractions from (Bi) were subjected to immunoprecipitations with rabbit anti-Bag6 antibody or rabbit control IgG antibody. Inputs and immunoprecipitates were analysed by immunoblotting for the indicated endogenous proteins. SGTA served as loading control as well as an internal control for equal Bag6 co-immunoprecipitation potential. (D) Mean ± s.e.m. of MAVS levels that co-immunoprecipitate with Bag6 in siATP13A1-treated cells normalised to nt siRNA-treated cells for four independent experiments as shown in (C). Same colour data points correspond to a single biological replicate; ns, not significant (paired two-tailed t test).

### EMC5 deficiency enhances BAG6-MAVS interaction

Prompted by the potential ER mislocalisation of a population of mitochondrial MAVS (McKenna et al., 2020), we next asked whether the association of MAVS with the BAG6 complex occurs before MAVS insertion into the ER membrane. Earlier work had already established that MAVS is integrated into cellular membranes via a TRC40-WRB/CAML-independent pathway (Coy-Vergara et al., 2019). We therefore focussed our attention on the EMC, which provides an alternative route for the membrane insertion of certain TA proteins (cf. (Coy-Vergara et al., 2019)) and shows structural homology to the CAML subunit of the TRC40-dependent membrane insertion pathway (McDowell et al., 2020). In this scenario, if MAVS accesses an EMC-mediated pathway for ER insertion operating downstream of BAG6, then any perturbation of this pathway may alter the pool of MAVS available to bind to BAG6 (Fig. 5A).

**Fig. 5.**
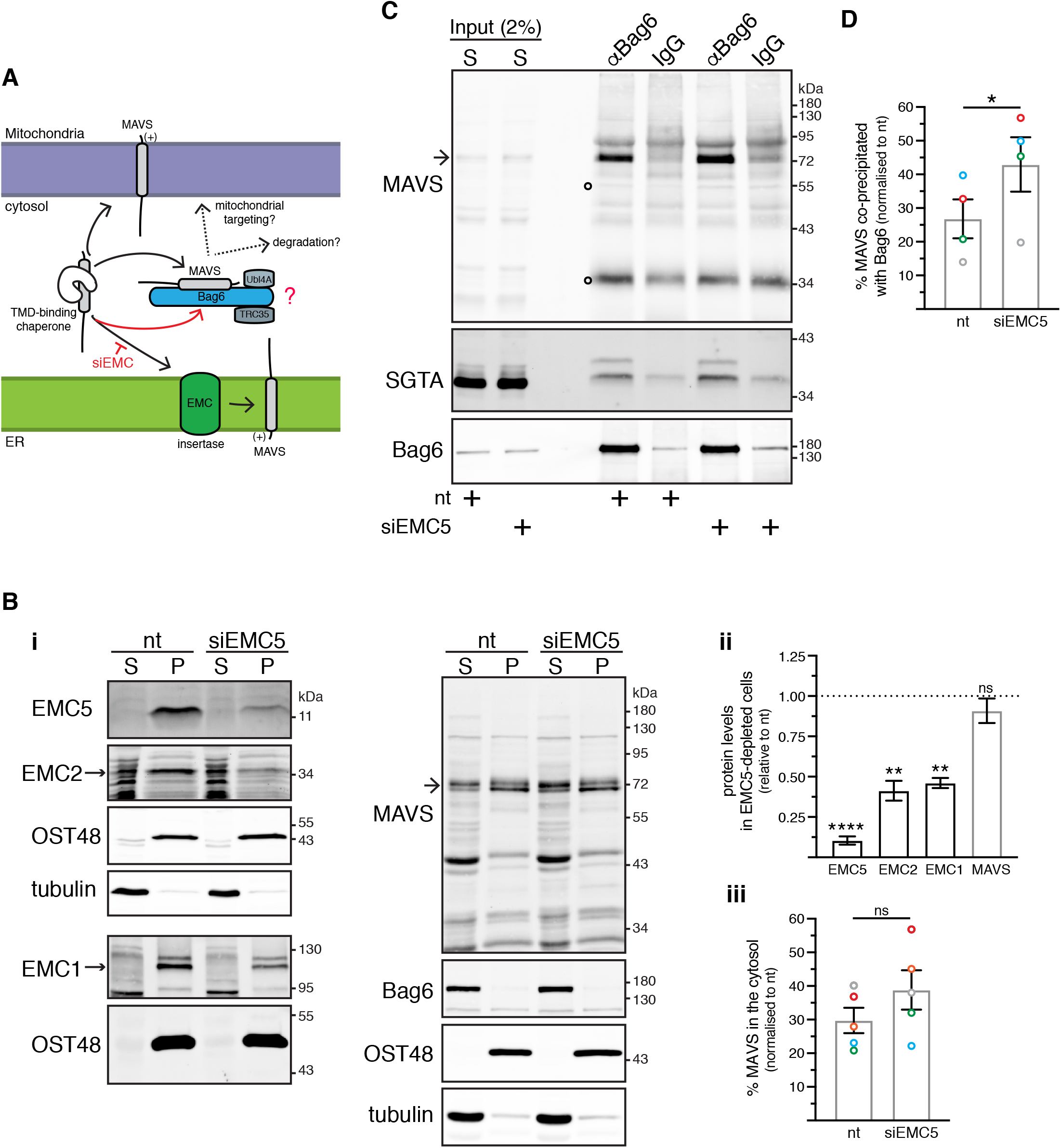
EMC5 depletion enhances Bag6-MAVS interaction. (A) Proposed model: Bag6 binds MAVS before its EMC-mediated integration at the ER membrane. EMC deficiency will promote Bag6 binding to the cytosolic pool of MAVS that fails to be imported into the ER membrane. (B) EMC5 depletion does not grossly alter the levels of MAVS in the crude cytosolic supernatant fraction. (i) Control KO cells transfected with non-targeting (nt) or EMC5-targeting siRNAs were fractionated as shown in Fig. 2A. Equivalent amounts of supernatant (S) and pellet (P) fractions were analysed by immunoblotting for the indicated endogenous proteins. (ii) EMC subunit and MAVS levels in siEMC5-treated cells relative to nt siRNA-treated cells, where protein levels were set to 1. Shown are means ± s.e.m. for three-five biological replicates as shown in (Bi). ****P < 0.0001; **P < 0.01; ns, not significant (one-way ANOVA with Tukey’s multiple comparison tests). (iii) Mean ± s.e.m. of the supernatant/total ratio of MAVS levels in siEMC5-treated cells normalised to nt siRNA-treated cells for five independent experiments as in (Bi). Same colour data points correspond to a single biological replicate; ns, not significant (paired two-tailed t test). (C) EMC5 depletion enhances Bag6-MAVS interaction. Supernatant (S) fractions from (Bi) were subjected to immunoprecipitations with rabbit anti-Bag6 antibody or rabbit control IgG antibody. Inputs and immunoprecipitates were analysed by immunoblotting for the indicated endogenous proteins. SGTA served as loading control as well as an internal control for equal Bag6 co-immunoprecipitation potential. (D) Mean ± s.e.m. of MAVS levels that co-immunoprecipitate with Bag6 in siEMC5-treated cells normalised to nt siRNA-treated cells for four independent experiments as shown in (C). Same colour data points correspond to a single biological replicate; * P < 0.005 (paired two-tailed t test).

To explore this possibility, we compared the cytosolic population of MAVS bound to BAG6 in both control and EMC-depleted cells by knocking down the ‘core’ EMC5 subunit (see also (O’Keefe et al., 2021b)). Notably, in HepG2 cells, we achieved a ∼90% depletion of EMC5 accompanied by a significant reduction of two other ‘core’ EMC subunits, EMC1 and EMC2 (Fig. 5Bi,ii), presumably destabilising the entire EMC (see also (O’Keefe et al., 2021b)). Although an EMC5 knockdown did not noticeably alter the distribution of MAVS following fractionation (Fig. 5Bi,ii), the BAG6-bound pool of MAVS was increased by ∼60% following EMC5 depletion when compared to control cells (Fig. 5C,D), consistent with a smaller, although non-significant increase (∼30%) in cytosolic MAVS (Fig. 5Bi,iii). These results are consistent with a model where the pool of BAG6-bound MAVS is dependent on the amount of cytosolic MAVS that is available for binding. Furthermore, since we see an effect of perturbing MAVS insertion into, but not its dislocation from, the ER membrane (cf. Figs 4, 5), we conclude that the BAG6 complex most likely captures newly synthesised MAVS before its membrane integration (see Discussion).

### BAG6-MAVS interaction is modulated during MAVS-dependent innate immune signalling

MAVS plays a central role in innate immune responses to the RNA virus infection of mammalian cells, acting downstream of viral RNA receptors which, upon binding MAVS at the MOM, trigger the formation of prion-like MAVS aggregates (Thoresen et al., 2021). These then serve as signalling platforms for the activation of interferon regulatory type 3 (IRF3) and nuclear factor kappa B (NF-κB) transcription factors (Thoresen et al., 2021) (Fig. S6A). Following activation, MAVS is ubiquitinated and subsequently targeted for proteasomal or lysosomal degradation, thereby diminishing downstream signalling (Ren et al., 2020).

Given the well-established role of the BAG6 complex in the quality control of hydrophobic precursor proteins that have mislocalised to the cytosol (Benarroch et al., 2019; Casson et al., 2017), we investigated the possibility that it may be involved in the regulated degradation of MAVS. To test this, we transfected control or Bag6- depleted cells with high-molecular weight poly(I:C), a non-viral ligand that is a specific activator of the RIG-I-like receptor MDA5 (Kato et al., 2008) (Fig. S6A), and examined the activation of IRF3, a signalling pathway component acting downstream of MAVS. No obvious effects on either the phosphorylation or dimerisation of IRF3 were detected following knockdown of the Bag6 protein (Fig. S6B,C), suggesting that the BAG6 complex is not an essential component for the cellular mobilisation of a MAVS-dependent response to viral infection.

We next addressed whether the activation of MAVS-dependent signalling might modulate the cytosolic pool of MAVS that is bound to the BAG6 complex. Stimulation with poly(I:C) induced phosphorylation and dimerisation of IRF3 but had no effect on total IRF3 levels (Fig. 6A). In agreement with the previously observed downregulation of MAVS expression following virus-induced activation of MAVS-mediated signalling (Ren et al., 2020), prolonged poly(I:C) stimulation resulted in a substantial ∼70% reduction in membrane-associated MAVS after 24 h (Fig. 6Bi, ii). In contrast, the relative amount of MAVS recovered in the cytosol was unaltered across the same 24 h time-course (Fig. 6Bi,iii). Interestingly, the proportion of this cytosolic pool of MAVS that was recovered with Bag6 initially showed a transient decline of ∼30% at 4 h and 12 h after stimulation, but returned to its initial level at 24 h (Fig. 6C,D). These data show that the population of MAVS that is bound to the BAG6 complex is dynamic and may respond to the activation of innate immune signalling following viral infection.

**Fig. 6.**
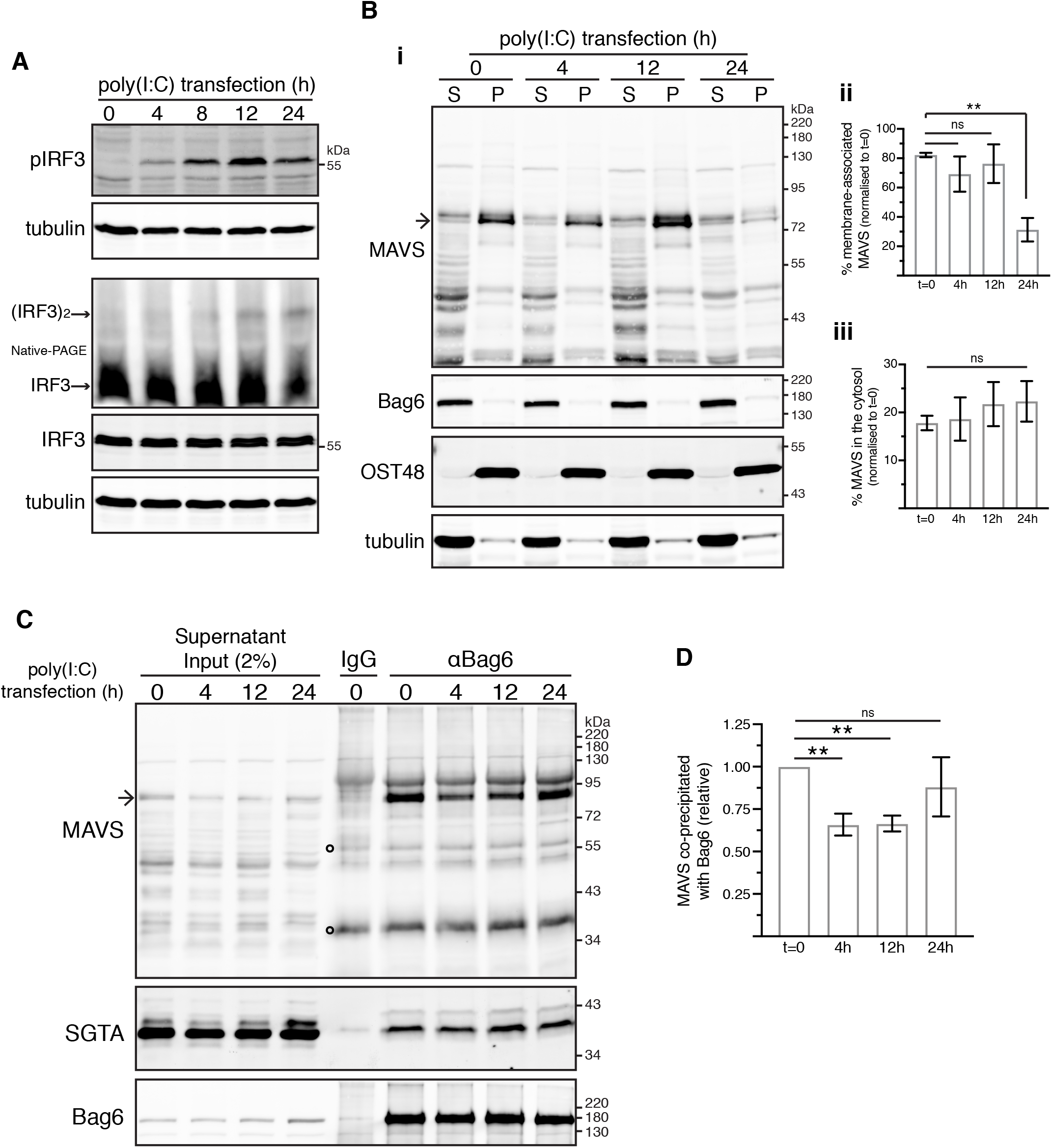
Stimulation with poly(I:C) compromises Bag6-MAVS interaction. (A) Kinetics of IRF3 activation in response to cytosolic poly(I:C). Control KO cells were mock- transfected (t=0) or transfected with poly(I:C) for various times before immunoblotting for the indicated proteins. Activation of endogenous IRF3 was assessed by induction of its phosphorylation and dimerisation. (B) Stimulation with cytosolic poly(I:C) does not grossly alter the levels of MAVS in the crude cytosolic supernatant fraction. (i) Control KO cells were mock-transfected (t=0) or transfected with poly(I:C) for various times before their fractionation as shown in Fig. 2A. The resulting supernatant (S) and pellet (P) fractions were analysed by immunoblotting for the indicated endogenous proteins. (ii, iii) Mean ± s.e.m. of the (ii) pellet/total ratio and (iii) supernatant/total ratio of MAVS levels in poly(I:C)-transfected cells normalised to mock-transfected cells (t=0) for six independent experiments as in (Bi). **P < 0.01; ns, not significant (ordinary one-way ANOVA with Dunnett’s multiple comparison tests). (C) Cytosolic poly(I:C) impairs Bag6-MAVS interaction. Supernatant fractions from (Bi) were subjected to immunoprecipitations with rabbit anti-Bag6 antibody or rabbit control IgG antibody. Inputs and immunoprecipitates were analysed by immunoblotting for the indicated endogenous proteins. SGTA served as loading control as well as internal control for comparable Bag6 binding. (D) Mean ± s.e.m. of MAVS levels that co-immunoprecipitate with Bag6 in poly(I:C)- transfected relative to mock-transfected cells (t=0) for six independent experiments as shown in (C). *P < 0.05; **P < 0.01; ns, not significant (one-way ANOVA with Dunnett’s multiple comparison tests).

## DISCUSSION

SGTA and the BAG6 complex are early TMD-recognition factors acting on the major pathway for TA targeting to the ER (Casson et al., 2017; Farkas and Bohnsack, 2021; Hegde and Keenan, 2021). In this study, we have identified the mitochondrial TA protein MAVS as an endogenous client of both SGTA and the BAG6 complex. The TMD region of MAVS is essential for its binding to SGTA, consistent with its dynamic engagement of newly synthesised TA proteins (Casson et al., 2017; Farkas and Bohnsack, 2021; Hegde and Keenan, 2021), including Stx-5 which is destined for the ER. Although MAVS is primarily localised to the MOM, it is also found at both MAMs (i.e. ER domains juxtaposed to discrete sites on mitochondria (Wu et al., 2018)) and peroxisomes (Horner et al., 2011). Whilst the combination of a moderately hydrophobic TMD and a positively charged C-terminal region (Fig. S4C) are necessary to reach these distinct locations (Dixit et al., 2010), it is unclear exactly how such diversity of targeting is achieved for MAVS.

Previous studies suggest that some newly synthesised TA proteins that are destined for mitochondria, such as OMP25, are bound by cytosolic Ubiquilins, which can prevent their aggregation prior to insertion into the MOM (Itakura et al., 2016). However, we were unable to find any evidence of a stable interaction between the Ubiquilins and OMP25 or MAVS in our experimental system (Fig. S5), leaving open their contribution to the biogenesis of TA proteins destined for the MOM (Fig. 7, pathway 1). In contrast to OMP25, we recover a pool of cytosolic MAVS bound to the BAG6 complex, most likely acting downstream of SGTA, which can capture nascent TA clients as their hydrophobic TMDs emerge from the ribosomal exit tunnel (Leznicki and High, 2020) (Fig. 7, pathway 2). In further support of this model, we find that the proportion of MAVS recovered with the BAG6 complex is reduced upon SGTA knockout (Fig. 3). Nevertheless, SGTA is dispensable for MAVS binding to the BAG6 complex, suggesting that there are alternative mechanisms for loading MAVS onto the BAG6 complex (Fig. 7, pathway 3).

**Fig. 7.**
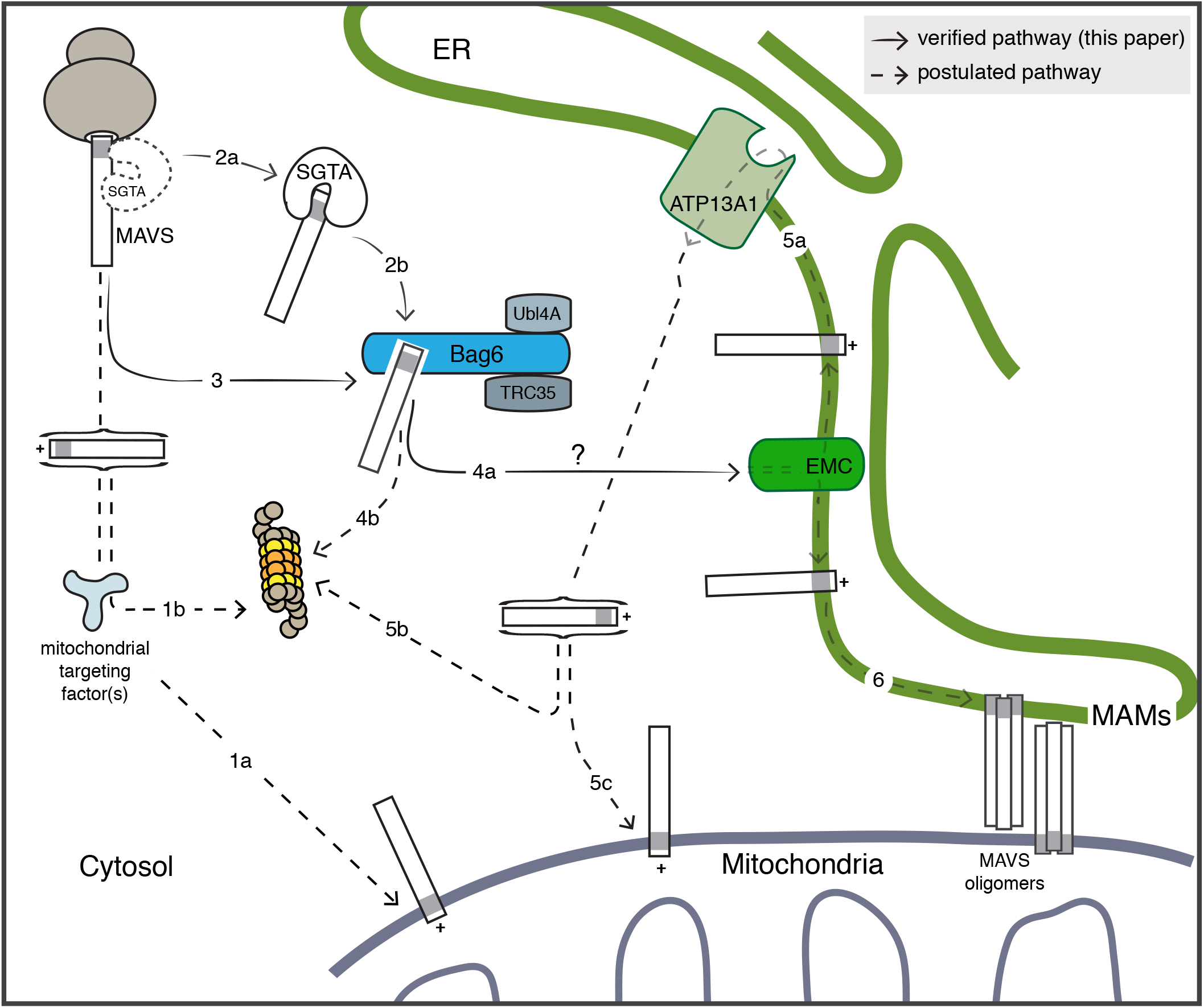
Working model for the role of the BAG6 complex during MAVS biogenesis. The molecular basis for the post-translational targeting and insertion of TA proteins such as MAVS into the mitochondrial outer membrane (MOM) are poorly defined (Bykov et al., 2020) (pathway 1a). Failed mitochondrial import can result in the mislocalisation of MAVS to the cytosol, where it may be recognised by one or more quality control machineries and targeted for proteasomal degradation (Itakura et al., 2016) (pathway 1b). A fraction of newly synthesised MAVS also engages the BAG6 complex either directly (pathway 3) or after transfer from SGTA (pathway 2). In the later case, SGTA may bind MAVS as its TMD leaves the ribosomal exit tunnel or after its release into the cytosol (Leznicki and High, 2020). The BAG6 complex acts upstream of MAVS ‘misinsertion’ into the ER membrane, which is most likely facilitated by the EMC (pathway 4a). It is currently unknown whether additional factors act between the BAG6 complex and the EMC insertase (pathway 4a, see ?). BAG6 binding might also enable the proteasomal degradation of mislocalised MAVS (Rodrigo-Brenni et al., 2014) (pathway 4b). At the ER membrane, ‘mistargeted’ MAVS can be recognised by the P5A-ATPase ATP13A1 and extracted to the cytosol (McKenna et al., 2020) (pathway 5a) for either proteasomal degradation (pathway 5b) or reinsertion into the MOM via an ER-SURF pathway (Hansen et al., 2018) (pathway 5c). The access of an ER-localised pool of MAVS to ER-MOM contacts sites (MAMs) that facilitate MAVS oligomerisation and downstream signalling (Esser-Nobis et al., 2020) (pathway 6) may be modulated by the innate immune response.

The selective binding of MAVS to the BAG6 complex in cytosolic extracts and the exogenous Bag6 protein expressed *in cellulo* (Fig. 3, cf. OMP25), most likely reflects a stable interaction between MAVS and the Bag6 subunit of the heterotrimeric BAG6 complex, which acts as a holdase that can also initiate the selective ubiquitination of aberrant clients such as MLPs (Leznicki et al., 2013; Rodrigo-Brenni et al., 2014; Wang et al., 2011). This stable binding of MAVS to the Bag6 subunit is distinct from the fate of SGTA-bound clients that are on a productive pathway for ER targeting (Casson et al., 2016). In this case, the Ubl4A subunit of the BAG6 complex facilitates a rapid and privileged transfer of these TA proteins from SGTA directly to TRC40 (Mock et al., 2015; Shao et al., 2017). However, although MAVS is detected in both the MOM and the ER (Esser-Nobis et al., 2020), it was neither bound by a TRC40 trap mutant nor affected by co-depletion of TRC40 and WRB (Coy-Vergara et al., 2019). Hence, in contrast to Stx-5 and other TA proteins that are efficiently targeted to the ER, the biogenesis of MAVS is TRC40-independent (Coy-Vergara et al., 2019).

Taken with our own findings, these data suggest that SGTA binds to MAVS but is unable to hand it off to TRC40, and therefore relies on alternative downstream acceptors for this client including the Bag6 subunit of the BAG6 complex. This model is in good agreement with the suggestion that the BAG6 complex acts as a sortase that directs hydrophobic MLPs towards either ER targeting or selective degradation, depending on the ability of TRC40 to receive SGTA-bound clients (Casson et al., 2016; Farkas and Bohnsack, 2021). Although a knockdown of the Bag6 protein has no clear effect on steady-state MAVS levels (Fig. S7), any contribution of the BAG6 complex to its proteasomal degradation (Fig. 7, pathway 4b) may be redundant (Rodrigo-Brenni et al., 2014). We therefore conclude that the BAG6 complex more likely acts as a “holdase” for mislocalised cytosolic MAVS that shields its hydrophobic TA region (Itakura et al., 2016; Wang et al., 2011). Whether BAG6-bound MAVS is maintained in a soluble conformation that is competent for subsequent membrane insertion and/or this specific pool of MAVS can be selectively ubiquitinated to enable its proteasomal degradation (Fig. 7, pathway 4) will require further detailed studies (see (Casson et al., 2016; Farkas and Bohnsack, 2021; Hegde and Keenan, 2021)).

In order to address at what stage of MAVS biogenesis it can interact with the BAG6 complex, we perturbed a defined pool of MAVS that mislocalises to the ER membrane (McKenna et al., 2020). When the P5-ATPase-dependent extraction of ER-localised MAVS was perturbed by knocking down ATP13A1 (McKenna et al., 2020), we found no reduction in the amount of MAVS that is recovered with the BAG6 complex (Fig. 4). However, when the EMC-dependent misinsertion of MAVS into the ER membrane was inhibited by knocking down the EMC5 subunit, a significant increase in the cytosolic pool of BAG6-bound MAVS was observed (Fig. 5). On this basis, we conclude that MAVS is engaged by BAG6 prior to its misinsertion into the ER membrane via an EMC-dependent pathway (Fig. 7, pathway 4) that likely reflects the structural conservation of the EMC and WRB/CAML complex (McDowell et al., 2020).

Whilst the delivery of mitochondrial TA proteins to the ER membrane is typically described as mislocalisation, it has now been established that a subset of membrane proteins can be productively redirected to the MOM from the ER via a process known as ER-SURF (Koch et al., 2021). It is currently unclear whether mitochondrial TA proteins, including MAVS and OMP25, are competent for onward delivery to the MOM following their ATP13A1-dependent extraction from the ER (Fig. 7, pathway 5) (see (Farkas and Bohnsack, 2021; McKenna et al., 2020)). Hence, it is possible that SGTA, the BAG6 complex and the EMC enable the ER insertion of MAVS proteins that are on a productive route to the MOM via the ER (Fig. 7, pathways 2-5). It should also be noted that a pool of MAVS, which localises to ER-MOM contact sites (MAMs), has been proposed to play a key role in initiating the innate immune response to viral infection (Esser-Nobis et al., 2020; Thoresen et al., 2021). The mechanism(s) by which such a MAM-localised pool of MAVS is generated remains undefined and we propose that one function of a BAG6/EMC-mediated pathway for ER insertion may be to provide a subpopulation of ER-localised MAVS that is competent for sorting into such ER-MOM contacts (Fig. 7, pathway 6). Hence, following insertion into the ER, the MAVS protein may have access to multiple fates that are regulated by prevailing cellular conditions including viral infection. In this context, when an artificial innate immune response that activates MAVS-dependent signalling is induced (Fig. 6A), we initially observe a significant reduction in the cytosolic pool of MAVS that is recovered with the BAG6 complex (Fig. 6). However, this pool of BAG6-bound MAVS returns to its pre-stimulation level after 24 h, reflecting a reduction in downstream IRF3 phosphorylation (Fig. S6A) that is observed over the same time frame (Fig. 6). Thus, the pool of BAG6-bound MAVS that we identify in this study appears to be subject to temporal regulation during the innate immune signalling process, perhaps allowing for the fine-tuning of MAVS availability in selected organellar membranes (see (Vazquez and Horner, 2015)) (Fig. 7). Further studies will be required to establish whether this BAG6-bound pool of MAVS plays any specific role in the propagation of RIG-I-like receptor-dependent signalling following viral infection (Thoresen et al., 2021).

## MATERIALS AND METHODS

### Antibodies and reagents

Anti-Bag6 rabbit polyclonal antibodies [1:1000 for immunoblotting (IB); 1:100 for immunoprecipitation (IP)] were raised against a synthetic peptide corresponding to residues 112-130 of human Bag6 isoform 2 (GSPPGTRGPGASVHDRNAN; synthesised by Peptide Specialty Laboratories GmbH) and affinity purified (also described in (Leznicki et al., 2013)). Anti-SGTA chicken polyclonal antibodies (1:2500 for IB; 1:200 for IP) were raised against recombinant His-thioredoxin-SGTA and affinity purified (also described in (Leznicki et al., 2015)). Other antibodies used were as follows: anti- ATP13A1 (Proteintech 16244-1-AP; 1:2500 for IB), anti-Bag6 (Abnova H00007917- B01P; 1:5000 for IB), anti-Calnexin (Cell Signaling Technology 2679S; 1:1000 for IB), anti-EMC1 (Abgent AP10226b; 1:500 for IB), anti-EMC2 (Santa Cruz Biotechnology sc- 166011; 1:500 for IB), anti-EMC5 (Bethyl Laboratories A305-832A-M; 1:1000 for IB), anti-Grasp65 [gift from Jon Lane, University of Bristol, UK; 1:5000 for immunofluorescence (IF)], anti-HA (Covance MMS-101R; 1:1000 for IB), anti-His (Sigma H1029; 1:3000 for IB), anti-Hsp70 (Abcam ab47455; 1:5000 for IB), anti-Hsp90 (Enzo Life Sciences ADI-SPA-846; 1:2000 for IB), anti-IRF3 (Cell Signaling Technology #11904; 1:1000 for IB), anti-pho-IRF3 (Cell Signaling Technology #4947; 1:1000 for IB), anti-MAVS (Santa Cruz Biotechnology sc-166583; 1:1000 for IB), anti-MAVS (Enzo Life Sciences ALX-210-929-C100; 1:50 for IF, 1:100 for IP), anti-Myc (Merck Millipore 05-724; 1:5000 for IB), anti-OMP25 (Proteintech 15666-1-AP; 1:1000 for IB, 1:50 for IF), anti-OST48 (previously described (Roboti and High, 2012); 1:1000 for IB), anti-SGTA (Santa Cruz Biotechnology sc-130557; 1:500 for IB), anti-Stx-5 (Synaptic Systems 110053; 1:5000 for IB, 1:50 for IF, 1:100 for IP), anti-Stx-5 (Santa Cruz Biotechnology sc-365124; 1:500 for IB), anti-TOM20 (Santa Cruz Biotechnology sc- 17764; 1:500 for IB, 1:50 for IF), anti-TRC35 (Bethyl Laboratories A302-613A; 1:1000 for IB), anti-tubulin (gift from Keith Gull, University of Oxford, UK; 1:1000 for IB), anti-Ubiquilin (Invitrogen 37-7700; 1:1000 for IB), anti-Ubiquilin-2 (Abcam ab217056; 1:2000 for IB, 1:100 for IP), anti-V5 (Novus Biologicals NB600-379; 1:1000 for IF), anti-rabbit IgG (Santa Cruz Biotechnology sc-2027; 1:40 for IP) and anti-chicken IgY (Santa Cruz Biotechnology sc-2718; 1:40 for IP). For infrared IB, IRDye 680LT/800CW- conjugated secondary antibodies (1:5000) raised in donkey were purchased from LI-COR BioSciences. For IF, Alexa Fluor 488-conjugated donkey anti-chicken (1:1000), Cy5-conjugated donkey anti-rabbit (1:400) and Cy3-conjugated donkey anti-mouse/sheep (1:1000) antibodies were purchased from Jackson ImmunoResearch Laboratories. Other commercially available reagents were: PMSF (Sigma-Aldrich 93482; 1 mM final), protease inhibitor cocktail (Sigma-Aldrich P8340; 1:100), phosphatase inhibitor cocktail set II (Calbiochem 524625; 1:100), benzonase nuclease (Millipore 70746; 250 U final) and EasyTag EXPRESS [^35^S]Met/Cys mix (PerkinElmer NEG772014MC).

### Plasmids and siRNA oligonucleotides

SGTA-V5, Bag6-V5 and PEX19-V5 in pcDNA5/FRT/V5-His-TOPO were previously described (Leznicki and High, 2012; Leznicki et al., 2015; Payapilly and High, 2014). pcDNA3.1-Myc-BioID2 (#74223) and pcDNA3.1-BioID2-HA (#74224) plasmids were obtained from Addgene. The sequences encoding SGTA-V5 and PEX19-V5 were amplified from pcDNA5 and inserted into the NheI-AgeI sites of pcDNA3.1-BioID2- HA. The plasmids encoding substrate_bd_mt and (UBL_bd_ & TPR_d_)mt SGTA-V5-BioID2-HA were generated by site-directed mutagenesis with PfuTurbo DNA polymerase (Agilent Technologies). Full-length human MAVS cDNA was transferred from pGEM- MAVS (Sino Biological HG15224-G) into the KpnI-NotI sites of the pCMV6-Entry vector (OriGene Technologies #PS100001) and an N-terminal FLAG tag was introduced by site-directed mutagenesis. The CARD (residues 10-77) and TMD (residues 514-535) were deleted from FLAG-MAVS coding sequence by inverse PCR. The siRNA oligonucleotides used were: ON-TARGETplus non-targeting control pool (Dharmacon D-001810-10-20), ON-TARGETplus human siATP13A1 (Dharmacon J- 020426-06-0020), siEMC5 (ThermoFisher Scientific s41129), siBag6#1 (CAGCUCCGGUCUGAUAUACAA; (Winnefeld et al., 2006)), siBag6#2 (UUUCUCCAAGAGCAGUUUA; (Minami et al., 2010)) and siBag6#3 (GCUCUAUGGCCCUUCCUCA). The siBag6 duplexes were made to order as ‘ON-TARGETplus’ by Dharmacon.

### Cell culture and transfection

Parental HepG2 cells (ATCC #85011430) and HepG2-derived KO cell lines were cultured in high-glucose DMEM (Sigma-Aldrich) supplemented with 10% (v/v) FBS (Sigma-Aldrich) at 37°C, 5% CO_2_ in a humidified incubator. Cells were routinely tested for mycoplasma contamination. DNA transfection was performed using Lipofectamine 3000 (ThermoFisher Scientific) by preparing a DNA solution containing P3000 reagent (1.5 μl/μg DNA), and then forming complexes of DNA:Lipofectamine 3000 at a ratio of 2:3. Transfection with poly(I:C) (Invivogen #tlrl-pic; 1 μg/ml final) was performed using Lipofectamine 3000 as described above. For siRNA transfection, cells were transfected with 25 nM siRNA oligonucleotides using Lipofectamine RNAiMAX (ThermoFisher Scientific) according to manufacturer’s instructions.

### Generation of SGTA KO cells

KO cell lines were generated using the CRISPR-Cas9 system as described (Ran et al., 2013). Briefly, two double-stranded oligonucleotides targeting exon 2 of human SGTA (guide #1: 5′-CATGACCAGCTCCGGCACGG-3′ and guide #2: 5′- CAGGAACTGGATGATGGCGT-3′) were ligated into the pSpCas9(BB)-2A-puro vector (Addgene #62988) using BbsI sites. HepG2 cells plated at 5 × 10^5^ cells per 3.5 cm dish were grown for 20 h before transfection with 2.5 μg of the resultant single-guide RNA (sgRNA) expression constructs using Lipofectamine 3000. Twenty-four hours post-transfection, transfected cells were selected by a 72 h incubation in media supplemented with puromycin 1 μg/ml. Puromycin-resistant cells were seeded into 96-well plates at 0.8 cells per well, and clonal cell lines screened for SGTA deficiency by IB and genomic sequencing.

### Immunoblotting

Denatured proteins were separated by SDS-PAGE and transferred to Immobilon-FL PVDF membranes (Merck Millipore IPFL00010). Membranes were incubated in a casein-based blocking buffer (Sigma-Aldrich) for 1 h and subsequently with primary antibodies in blocking buffer supplemented with 0.1% (v/v) Tween-20 for >12 h at 4°C. After three washes with TBST, membranes were incubated with secondary antibodies in blocking buffer supplemented with 0.1% (v/v) Tween-20 and 0.01% (v/v) SDS for 1 h at room temperature. For detecting biotinylated proteins, blocked membranes were incubated with IRDye800CW-conjugated streptavidin (LI-COR BioSciences 926-32230; 1:5000) for 1 h at room temperature. Membranes were washed extensively with TBST and fluorescent bands visualised on an Odyssey CLx Infrared Imager (LI-COR Biosciences). Signals were quantified using the automatic background subtraction function (setting: average, top and bottom) of the Image Studio Lite 5.2.5 software provided by the manufacturer.

### BioID2 labelling and streptavidin-affinity purification

A 2.05 mM biotin (Sigma-Aldrich B4501) stock was prepared in distilled water by brief sonication. Cells were seeded into 15 cm dishes at 7.5 × 10^6^ cells per dish and grown for ∼20 h before transfection with 30 μg of the indicated BioID2-tagged expression construct. About 36 h post-transfection, cells were incubated with fresh media supplemented with 50 μM biotin and harvested 8 h later. Cell pellets from two 15 cm dishes were solubilised in 1 ml RIPA buffer [50 mM Tris-HCl pH 7.5, 150 mM NaCl, 1 mM EDTA, 1 mM EGTA, 1% (v/v) IGEPAL CA-630, 0.5% (w/v) sodium deoxycholate and 0.1% (v/v) SDS] supplemented with PMSF, protease inhibitors and benzonase by continuous shaking at 4°C for 1 h. The lysates were sonicated (3 × 10 s bursts with 5 s rest) on ice at low amplitude in a Bioruptor (Diagenode) and then centrifuged at 20,817 × *g* for 30 min at 4°C. The post-nuclear supernatants were further diluted in RIPA buffer to 2.5 mg total protein per ml, and biotinylated proteins were affinity-purified by incubation with streptavidin-sepharose beads (Cytiva 17-5113-01; 8 μl 1:1 slurry per mg of total protein) at 4°C on a nutator for 3 h. The beads were pelleted (400 × *g*, 5 min) and washed four times with RIPA buffer and three times with ammonium bicarbonate 50 mM pH 8.0. After all residual ammonium bicarbonate was pipetted off, beads were flash-frozen and stored at - 80°C before shipping to the Proteomics Core facility at Sanford-Burnham-Prebys Medical Discovery Institute.

### On-beads protein digestion and LC-MS/MS analysis

Liquid-free beads were resuspended in 8 M urea dissolved in 50 mM ammonium bicarbonate and disulphide bonds were reduced with 10 mM tris(2- carboxyethyl)phosphine at 30°C for 60 min. After cooling the samples to room temperature, free cysteines were alkylated with 30 mM iodoacetamide for 30 min in the dark. Following alkylation, urea was diluted to 1 M using 50 mM ammonium bicarbonate, and proteins were subjected to overnight digestion with Mass Spec Grade Trypsin/Lys-C mix (Promega). The beads were then pulled down and the solutions containing the digested peptides were desalted using AssayMap C18 cartridges mounted on an AssayMap Bravo liquid handling system (Agilent Technologies) and subsequently dried down in a SpeedVac concentrator.

Prior to LC-MS/MS analysis, dried peptides were reconstituted in 2% (v/v) acetonitrile, 0.1% (v/v) formic acid and concentration determined using a NanoDrop spectrophotometer (ThermoFisher Scientific). Samples were then analyzed by LC-MS/MS using a Proxeon EASY-nanoLC system (ThermoFisher Scientific) coupled to a Q Exactive Plus Orbitrap mass spectrometer (ThermoFisher Scientific). Peptides were resolved on a 250 mm × 75 μm Aurora C18 reversed-phase analytical column (IonOpticks) over a 120 min organic gradient (1-5% solvent B over 1 min, 5-23% solvent B over 72 min, 23-34% solvent B over 45 min and 34-48% solvent B over 2 min) with a flow rate of 300 nl/min (60°C). Solvent A was 0.1% formic acid and solvent B was 80% acetonitrile in 0.1% formic acid. The mass spectrometer was operated in positive data-dependent acquisition mode. MS1 spectra were measured in the Orbitrap with a resolution of 70,000 (at m/z 400) in the mass range m/z 350- 1700. Automatic gain control (AGC) target was set to 1 x 10^6^ with a maximum injection time of 100 ms. Up to twelve MS2 spectra per duty cycle were triggered, fragmented by HCD, and acquired at a resolution of 17,500 and an AGC target of 5 x 10^4^, an isolation window of 1.6 m/z and a normalized collision energy of 25. The dynamic exclusion was set to 20 s with a 10 ppm mass tolerance around the precursor.

### MS data analysis

Raw data were analysed using MaxQuant software (v1.5.5.1) searching against the Uniprot *Homo sapiens* database (downloaded in January 2019) and the GPM cRAP database containing common contaminants. Precursor mass tolerance was set to 20 ppm for the first search, where initial mass recalibration was completed, and to 4.5 ppm for the main search. Product ions were searched with a mass tolerance of 0.5 Da. The maximum precursor ion charge state used for searching was 7. Cysteine carbamidomethylation was set as a fixed modification, while oxidation of methionine and acetylation of protein N-terminus were set as variable modifications. Enzyme was set to trypsin in a specific mode and a maximum of two missed cleavages was allowed for searching. The target-decoy-based false discovery rate (FDR) filter for spectrum and protein identification was set to 0.01. Protein label-free quantification (LFQ) intensities were exported from MaxQuant and analysed through SAINTexpress software (v3.6.3) (Teo et al., 2014) using default parameters to identify proximal interactions. Controls were set as both myc-BioID2 and PEX19-BioID2 samples. High-confidence bait-prey interactions were identified using a BFDR (Bayesian FDR) threshold of 0.05.

### Immunofluorescence analysis

Cells plated on coverslips at 5 × 10^5^ cells per 3.5 cm dish were transfected with 2.5 μg of the indicated plasmid using Lipofectamine 3000. Thirty-six hours post-transfection, cells were fixed with 4% (v/v) PFA in PBS for 30 min at room temperature and unreacted aldehyde groups quenched with 0.1 M glycine-Tris pH 8.5 for 12 min. Cells were then permeabilised with 0.1% (v/v) TX-100 for 5 min at room temperature and washed twice prior to incubation with primary antibodies in PBS for 1 h. Three washes with PBS followed before a further 1 h incubation with secondary antibody solution supplemented with the DNA dye DAPI. After cells were washed with PBS, coverslips were dried and mounted on slides with ProLong Gold Antifade reagent (ThermoFisher Scientific). Imaging was performed using an Olympus BX60 upright microscope using a 100 × oil-immersion objective and equipped with a MicroMax cooled, slow-scan CCD camera (Roper Scientific) driven by Metaview software (University Imaging Corporation). Images were processed with ImageJ.

### Cell fractionation and co-immunoprecipitation analysis

Three million cells were plated per 15 cm dish and harvested after ∼96 h. For cytosolic delivery of poly(I:C), cells were transfected 72 h after plating and harvested 4-24 h later. For knockdown experiments, cells plated at 7.5 × 10^6^ cells per 15 cm dish were transfected 20 h after plating and harvested 72 h post-transfection. Cell pellets were resuspended in 5 × volumes of buffer A (50 mM Tris-HCl pH 7.5, 150 mM NaCl and 1.5 mM MgCl_2_) supplemented with PMSF, protease and phosphatase inhibitors. The cell suspension was kept on ice for 30 min and then homogenised by 22 passes through a cell homogenizer (Isobiotec, Germany) with a tungsten carbide ball (clearance of 10 μm). The crude homogenate was subjected to two rounds of centrifugation at 700 × *g* for 10 min at 4°C and the resulting post-nuclear supernatant was further centrifuged at 21,380 × *g* for 30 min at 4°C to obtain a supernatant (crude cytosol) and a pellet (crude membranes) fraction. The supernatant fraction and the buffer A-washed pellet fraction were centrifuged again at 21,380 × *g* for 30 min at 4°C to remove contaminants. The supernatant fraction was then incubated with 5 μg of the indicated antibodies and protein A-agarose beads (GenScript L00210) at 4°C on a nutator for 5 h. After washing the beads four times with buffer A, bound proteins were eluted with 2 × SDS sample buffer and analysed by infrared IB.

### Pulse-chase analysis of protein secretion

The secretion assay was performed using a pulse-chase approach as previously described (Roboti et al., 2013). Briefly, cells were left untreated or treated with 2.5 μg/ml brefeldin A (Sigma-Aldrich B7651) for 1 h before being starved in Met- and Cys-free DMEM (ThermoFisher Scientific 21013024) for 20 min at 37°C. Labelling was initiated by addition of fresh starvation medium containing 22 mCi/ml [^35^S]Met/Cys for 15 min. After washing with PBS, cells were incubated in serum-free DMEM supplemented with 2 mM unlabelled L-Met and L-Cys for 2 h. Brefeldin A was included throughout the starvation, pulse and chase. Secreted proteins in the media were recovered by precipitation with 13% trichloroacetic acid, washed in acetone and dissolved in 2 × SDS sample buffer. Cells were solubilised with Triton X-100 lysis buffer (20 mM Tris pH 7.5, 150 mM NaCl, 1% (v/v) TX-100, 1 mM EDTA and 1 mM EGTA) supplemented with PMSF and protease inhibitors. Equal amounts of precipitated proteins in the media and clarified lysates were analysed by SDS-PAGE and phosphorimaging using FLA-3000 (Fujifilm).

### *In vitro* transcription, translation and pull-down assay

Linear DNA templates were generated by PCR using appropriate primers and transcribed into mRNA with T7 polymerase (Promega). *In vitro* translation was performed in rabbit reticulocyte lysate (Promega) supplemented with 1 mCi/ml [^35^S]Met/Cys, amino acid mix lacking Met (Promega) and 0.1 × volume of an *in vitro* transcribed mRNA (200-1,000 ng/μL) in the absence or presence of 2 μM recombinant His-S-tag-SGTA (gift from Pawel Leznicki, Sygnature Discovery, UK) for 90 min at 30°C. Following a 10 min incubation with 1 mM puromycin (Sigma-Aldrich P7255) at 37°C, the reactions were diluted 5.5-fold with buffer B (50 mM HEPES-KOH pH 7.5, 300 mM NaCl, 10 mM imidazole and 10% (v/v) glycerol) and incubated with pre-equilibrated HisPur Cobalt resin (ThermoFisher Scientific) at 4°C for 2 h. After washing the beads, bound proteins were eluted with buffer B supplemented with 200 mM imidazole. Samples were resolved by SDS-PAGE and radiolabelled products visualised by phosphorimaging using FLA-3000 (Fujifilm).

### Analysis of dimeric and phosphorylated state of IRF3

Native PAGE gel 7.5% was pre-run with running buffer [25 mM Tris-HCl pH 8.4 and 192 mM glycine in the presence of or absence of 0.5% (w/v) sodium deoxycholate in the cathode and anode buffer, respectively] at 120 V in a cold room for ∼35 min. Cells from confluent 3.5 cm dishes were lysed with ice-cold buffer A supplemented with 0.5% (v/v) IGEPAL CA-630, PMSF, protease and phosphatase inhibitors. The cell extracts were kept on ice for 30 min before centrifugation at 21,380 × *g* for 10 min to remove the insoluble fraction. The clarified cell lysates were then mixed with 5 × native sample buffer (250 mM Tris-HCl pH 6.8, 1% (w/v) sodium deoxycholate, 50% (v/v) glycerol and 0.5% (w/v) bromphenol blue) to a final concentration of 1 ×, and immediately loaded on the gel and run at 120 V for 100 min in a cold room. The monomeric and dimerised IRF3 were detected by infrared IB.

For detection of phosphorylated IRF3, cells from confluent 3.5 cm dishes were solubilised in ice-cold RIPA buffer supplemented with PMSF, benzonase, protease and phosphatase inhibitors by continuous shaking at 4°C for 1 h. The lysates were sonicated (3 × 10 s bursts with 5 s rest) on ice at low amplitude in a Bioruptor (Diagenode) and then centrifuged at 20,817 × *g* for 30 min at 4°C. Clarified lysates (∼150 μg total protein) were resolved by SDS-PAGE and analysed by infrared IB. Note that incubation of blots with anti-pho-IRF3 was performed in 5% BSA in TBST overnight at 4°C.

### Detergent solubility assay

The solubility assay was performed as described (Itakura et al., 2016) with minor modifications. Cells were lysed with ice-cold, non-denaturing lysis buffer [50 mM Tris-HCl pH. 7.4, 150 mM NaCl, 1mM EDTA, 0.5% (v/v) IGEPAL CA-630 and 0.5% (w/v) sodium deoxycholate] supplemented with PMSF and protease inhibitors by gentle shaking at 4°C for 30 min. After centrifugation at 20,817 × *g* for 30 min at 4°C, the post-nuclear supernatants (soluble fractions) were collected in a separate tube and the pellets (insoluble fractions) were dissolved in an equal volume of ice-cold RIPA buffer supplemented with PMSF, protease inhibitors and benzonase by vigorous shaking at 4°C for 1 h. Following a brief sonication step of the insoluble fractions on ice, both the soluble and the insoluble fractions were denatured in SDS sample buffer and analysed by infrared IB.

### Statistics

All experiments were repeated at least three times with representative data shown. Data are expressed as the mean ± s.e.m. and significant differences among two or multiple experimental groups were assessed by two-tailed t-test and one-way ANOVA, respectively. Statistical analyses and data plotting were performed using GraphPad Prism 9.

## ACKNOWLEDGEMENTS

We are grateful to Alexandre Rosa Campos and members of the Sanford Burnham Prebys Proteomics Core for their assistance with MS analysis. We also thank Martin Pool (The University of Manchester, UK) and Pawel Leznicki (Sygnature Discovery, UK) for their helpful insights and suggestions during the preparation of this manuscript.

