## Supplementary Figures for "Mitochondrial antiviral-signalling protein is a client of the BAG6 protein quality control complex"

### Figure S1

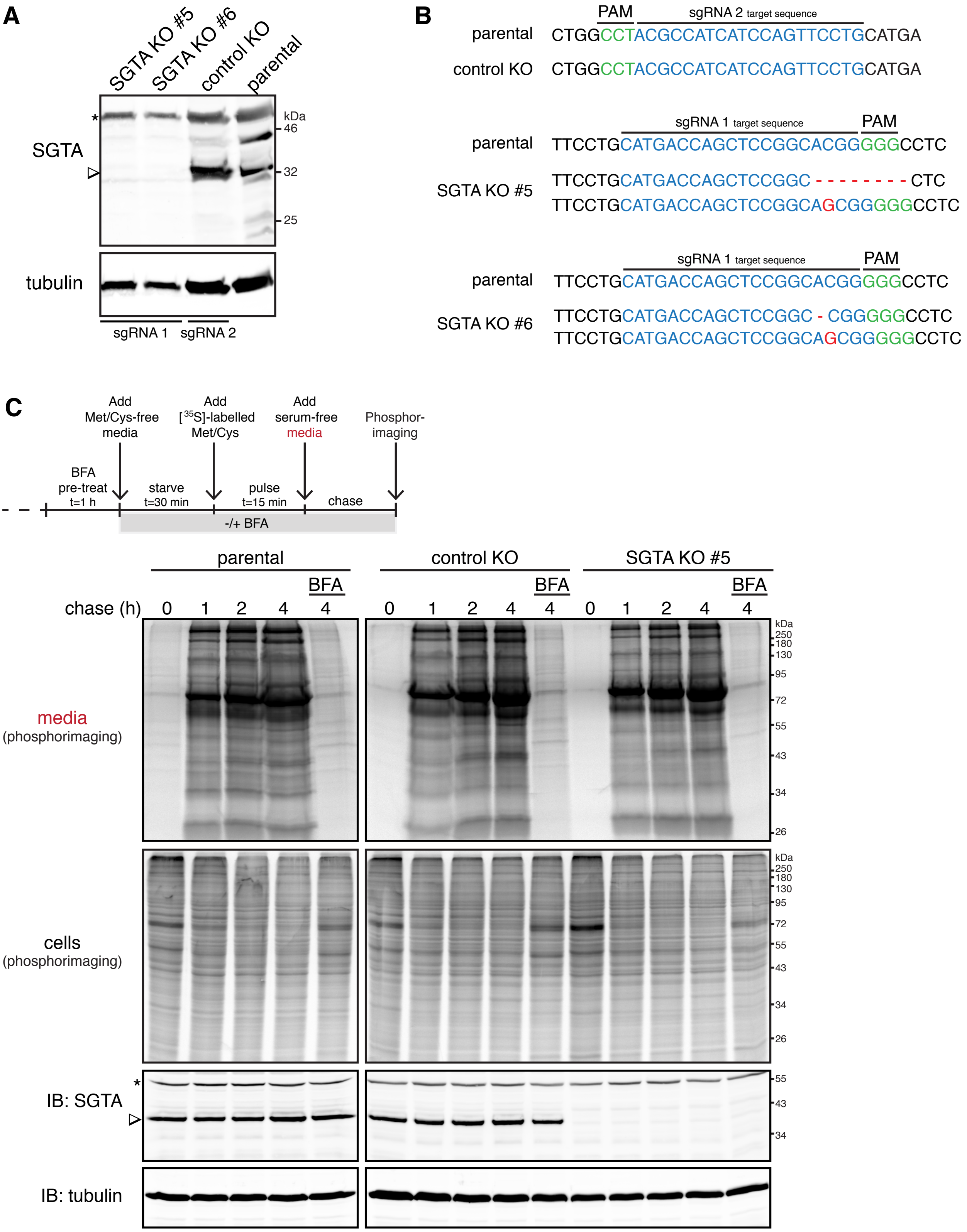

A
SGTA KO #5
SGTA KO #6
control KO
parental
SGTA
tubulin
kDa
46
32
25
sgRNA 1
sgRNA 2
B
PAM
sgRNA 2 target sequence
parental CTGGCCTACGCCATCATCCAGTTCCTGCATGA
control KO CTGGCCTACGCCATCATCCAGTTCCTGCATGA
sgRNA 1 target sequence
parental TTCCTGCATGACCAGCTCCGGCACGGGGGCCTC
SGTA KO #5
TTCCTGCATGACCAGCTCCGGC - - - - - - - - CTC
TTCCTGCATGACCAGCTCCGGCAGCGGGGGCCTC
SGTA KO #6
TTCCTGCATGACCAGCTCCGGC - CGGGGGCCTC
TTCCTGCATGACCAGCTCCGGCAGCGGGGGCCTC
C
Add Met/Cys-free media
Add [35S]-labelled Met/Cys
Add serum-free media
Phosphor-imaging
BFA pre-treat t=1 h
starve t=30 min
pulse t=15 min
chase
-/+ BFA
parental
control KO
SGTA KO #5
BFA
chase (h)
0 1 2 4 4
media (phosphorimaging)
cells (phosphorimaging)
IB: SGTA
IB: tubulin
250
180
130
95
72
55
43
34
26

**Fig. S1. Confirmation of SGTA KO in HepG2 cells.** CRISPR-Cas9-mediated disruption of exon 2 of the SGTA gene in HepG2 cells resulted in a loss of SGTA. For the generation of SGTA KO cells, we designed single guide RNAs (sgRNAs) targeting the exon 2 of human SGTA (see Materials and Methods), which is present in all putative SGTA isoforms (Ensemble accession ENSG00000104969). SGTA KO clonal cell lines #5 (used hereafter) and #6 were obtained using sgRNA 1. Control KO cells are clones that underwent the same manipulation with SGTA KO cells using sgRNA 2, but retained SGTA expression. (A) Total lysates from parental, control KO and SGTA KO HepG2 cells were analysed by immunoblotting with anti-SGTA and anti-tubulin antibodies. The SGTA signal is marked by an arrowhead, while a non-specific band retained in SGTA KO cells is indicated by an asterisk. (B) Sequence alignment of partial SGTA coding sequences from parental and control KO/SGTA KO HepG2 cells. PAM (protospacer adjacent motif), green; sgRNA target sequence, blue; mutated sequence, red. (C) SGTA deficiency does not inhibit global protein secretion. HepG2 cells were left untreated or treated with brefeldin A (BFA; 2.5 μg/ml) for 1 h. Newly synthesised proteins were pulse-labelled for 15 min and then chased in serum-free medium containing unlabelled Met and Cys for various times in the absence or presence of BFA, as shown in the schematic representation of the assay. At the end of the chase, radiolabelled proteins in media (upper panel) and whole cell lysates (bottom panel) were resolved by SDS-PAGE and visualised by phosphorimaging. Whole cell lysates were also analysed by immunoblotting (IB) with anti-SGTA and anti-tubulin antibodies. The arrowhead points to the SGTA signal and the asterisk indicates a non-specific band.

### Figure S2

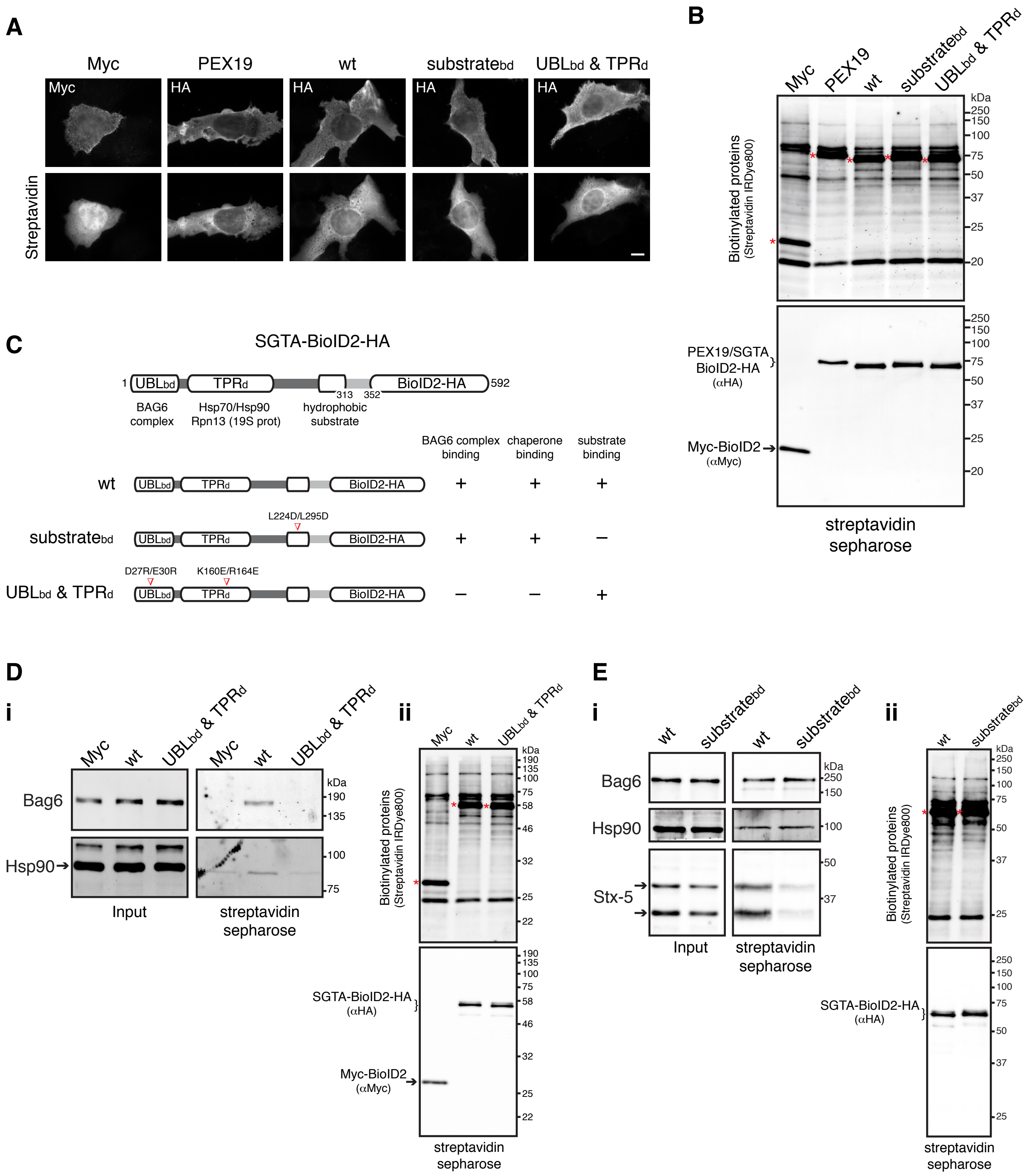

A
Myc
PEX19
wt
substratebd
UBLbd & TPRd
HA
Streptavidin
B
Biotinylated proteins
(Streptavidin IRDye800)
kDa
250
150
100
75
50
37
25
20
PEX19/SGTA
BioID2-HA
(αHA)
Myc-BioID2
(αMyc)
streptavidin
sepharose
C
SGTA-BioID2-HA
UBLbd
TPRd
BioID2-HA
1
313
352
592
BAG6
complex
Hsp70/Hsp90
Rpn13 (19S prot)
hydrophobic
substrate
BAG6 complex
binding
chaperone
binding
substrate
binding
L224D/L295D
D27R/E30R
K160E/R164E
D
i
ii
Bag6
Hsp90
Input
190
135
58
46
32
22
SGTA-BioID2-HA
E
Stx-5

**Fig. S2. Validation of BioID2-based assay for proximity labelling of SGTA proteome.** (A,B) Demonstration of the cytosolic localisation and the biotinylation activity of bait-BioID2 fusion proteins. SGTA KO cells transiently transfected with plasmids encoding Myc-BioID2, PEX19-BioID2 or the indicated SGTA-BioID2 variants were incubated with exogenous biotin for 8 h. (A) Cells were fixed and stained with anti-Myc/anti-HA and fluorophore-conjugated streptavidin to visualise BioID2-tagged baits and biotinylated proteins, respectively. Representative wide-field fluorescence images are shown. Scale bar, 10 μm. (B) Cells were lysed and biotinylated proteins were isolated on streptavidin sepharose beads. The streptavidin-bound material was eluted using a biotin-containing buffer, resolved by SDS-PAGE and immunoblotted for BioID2-tagged baits (anti-Myc/anti-HA) or probed with streptavidin to detect total biotinylated proteins in each sample. The signals from self-biotinylated BioID2-tagged baits were detected at the predicted molecular weights and are marked with red asterisks. (C-E) Detection of known SGTA interactors by a small-scale streptavidin pull-down assay. SGTA KO cells transiently expressing the indicated BioID2-tagged baits (see schematic in C; also shown in Fig. 1A) were treated with biotin for 8 h. (Di,Ei) Whole cell lysates (Input) and biotinylated proteins (streptavidin sepharose) were analysed by immunoblotting for the indicated endogenous proteins. Note that samples were analysed on two different gels and the combined image was produced using the same brightness and contrast settings. (Dii,Eii) Biotinylated proteins (streptavidin sepharose) were analysed by immunoblotting for BioID2-tagged baits (anti-Myc/anti-HA) and total biotinylated proteins (Streptavidin IRDye800). The signals from self-biotinylated BioID2-tagged baits are marked with red asterisks. Syntaxin-5 (Stx-5) appears as two bands (hereafter indicated by arrows) corresponding to two isoforms, a 42 kDa-ER and a 35 kDa-Golgi isoform that result from an alternative initiation of translation (Coy-Vergara et al., 2019).

### Figure S3

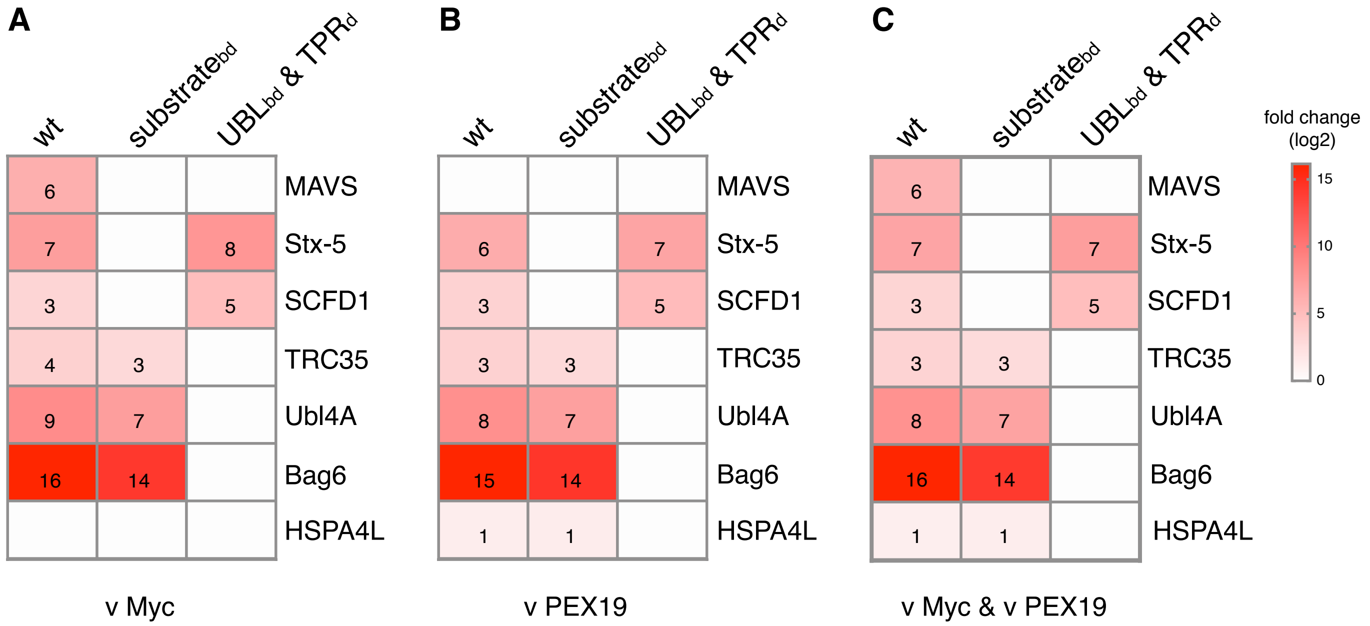
A
B
C
wt
substrate bd
UBL bd & TPR d
MAVS
Stx-5
SCFD1
TRC35
Ubl4A
Bag6
HSPA4L
6
7
8
3
5
4
3
9
7
16
14
6
7
3
5
3
3
8
7
15
14
1
1
6
7
7
3
5
3
3
8
7
16
14
1
1
v Myc
v PEX19
v Myc & v PEX19
fold change (log2)
15
10
5
0

**Fig. S3. Profiling of SGTA interactors by proximity-labelling proteomics.** Heat maps representing log2-transformed fold changes in the protein intensities of high-confidence (BFDR<0.05) wild-type (wt)/mutant SGTA-specific preys relative to (A) Myc-BioID2, (B) PEX19-BioID2 and (C) both Myc-BioID2 and PEX19-BioID2 controls. Individual rounded values are depicted in the heat maps. A non-significant prey is shown as a white box (three biological replicates, see Table S1 for list of all the detected proteins).

### Figure S4

A

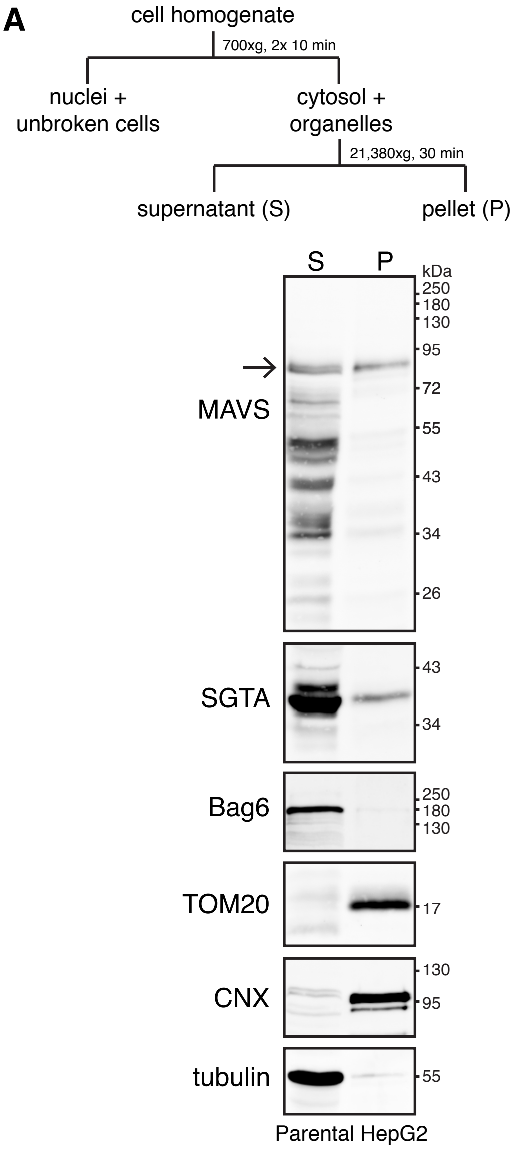


B

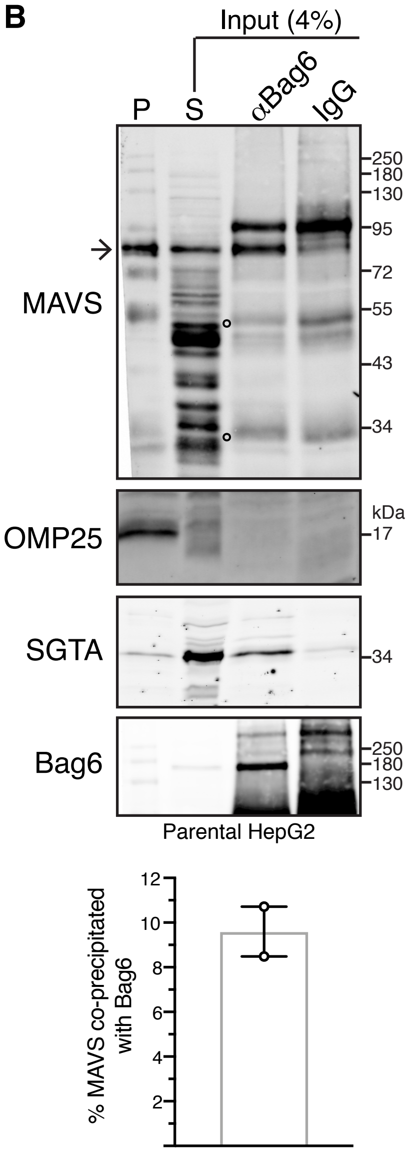


C

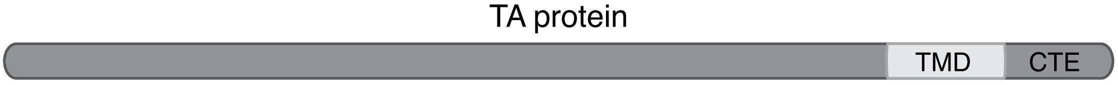


| TA protein | TMD | CTE | Predicted hydropathy | Predicted α-helicity | Destination membrane |
|---|---|---|---|---|---|
| MAVS | GALWLQVAVTGVLVVTLLVVL | YRRRLH | 2.34 | 0.18 | MOM / Peroxisomes |
| OMP25 | IPIFMVLVPVFALTMVAAWAF | MRYRQQL | 2.29 | 0.56 | MOM |
| Stx-5 | WLMVKIFLILIVFFIIFVVFL | A | 3.12 | 4.58 | ER |

**Fig. S4. Bag6-MAVS interaction in parental HepG2 cells.** (A) Top: Schematic of the subcellular fractionation protocol used to separate the cell homogenate into crude cytosolic supernatant (S) and membrane-associated pellet (P) fractions. Bottom: Detergent-free extracts from parental HepG2 cells were fractionated as shown above. Equivalent amounts of each fraction were analysed by immunoblotting for MAVS and various compartmental markers. Bag6, SGTA and tubulin (cytosolic markers), TOM20 (mitochondrial outer membrane marker) and Calnexin (CNX, ER membrane marker) serve as fractionation controls. Full-length MAVS species (hereafter marked by an arrow) can be observed in both the cytosolic and membrane-associated fractions at steady-state (see also Fig. 2A). (B) MAVS, but not OMP25, co-immunoprecipitates with Bag6. The supernatant fraction from (A) was subjected to immunoprecipitations with equal amounts of rabbit anti-Bag6 antibody or rabbit IgG antibody (control for non-specific binding). Equivalent amounts of pellet (P) and supernatant (S) fractions from (A) and immunoprecipitates were analysed by immunoblotting for the indicated endogenous proteins. SGTA served as a positive control for Bag6 binding. Open circles on MAVS blot indicate signals derived from denatured antibody heavy and light chains. In contrast to full-length MAVS that is distributed in both fractions, OMP25 is predominantly present in the membrane-associated pellet fraction. The levels of MAVS that co-immunoprecipitate with Bag6 were reported as mean ± s.e.m. for two independent experiments. (C) Sequences and properties of the transmembrane domains (TMDs) and C-terminal elements (CTEs) of the TA substrates used in this study. Sequences of TMDs were obtained from UniProt. For predicted hydropathy, Grand Average of Hydropathy (GRAVY) scores were calculated (http://www.gravy-calculator.de). For predicted $\alpha$-helicity, scores (% helical content) was calculated at pH 7.0, 310.15 K, ionic strength 0.13 M using the Agadir prediction algorithm (http://agadir.crg.es) (Munoz and Serrano, 1997). MOM, mitochondrial outer membrane; ER, endoplasmic reticulum.

### Figure S5

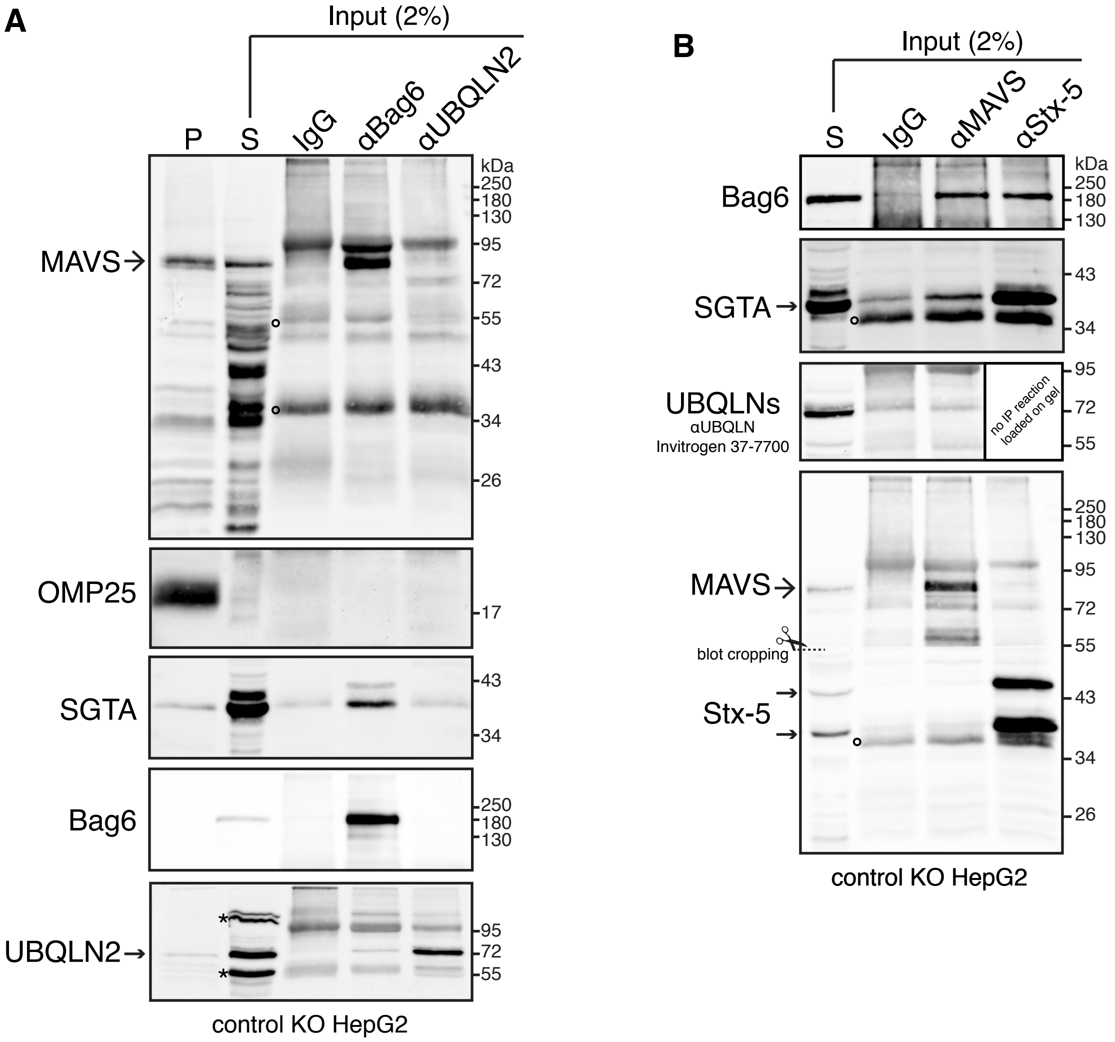
A
Input (2%)
P
S
IgG
αBag6
αUBQLN2
kDa
250
180
130
95
72
55
43
34
26
17
MAVS
OMP25
SGTA
Bag6
UBQLN2
*
control KO HepG2
B
Input (2%)
S
IgG
αMAVS
αStx-5
Bag6
SGTA
UBQLNs
αUBQLN
Invitrogen 37-7700
no IP reaction loaded on gel
MAVS
blot cropping
Stx-5
control KO HepG2

**Fig. S5. No association of MAVS with Ubiquilins can be detected by co-immunoprecipitation studies.** (A) MAVS does not co-immunoprecipitate with Ubiquilin-2 (UBQLN2). Control KO cells were fractionated as shown in the schematic of Fig. S4A and the supernatant fraction was subjected to immunoprecipitations with equal amounts of rabbit anti-Bag6, rabbit anti-UBQLN2 or rabbit IgG (control for non-specific binding) antibodies. Equivalent amounts of pellet (P) and supernatant (S) fractions and immunoprecipitates were analysed by immunoblotting for the indicated endogenous proteins. SGTA served as positive control for Bag6 binding. Asterisks indicate non-specific products. (B) Ubiquilins (UBQLNs) do not co-immunoprecipitate with MAVS. Control KO cells were fractionated as shown in the schematic of Fig. S4A and the supernatant (S) fraction was subjected to immunoprecipitations with equal amounts of rabbit anti-MAVS, rabbit anti-Stx-5 or rabbit IgG (control for non-specific binding) antibodies. Input and immunoprecipitates were analysed by immunoblotting for the indicated endogenous proteins. Note that the Invitrogen 37-7700 anti-UBQLN antibody recognises human UBQLN-1, -2 and -4 (Safren et al., 2015). In (A) and (B), open circles indicate signals derived from denatured antibody heavy and light chains.

### Figure S6

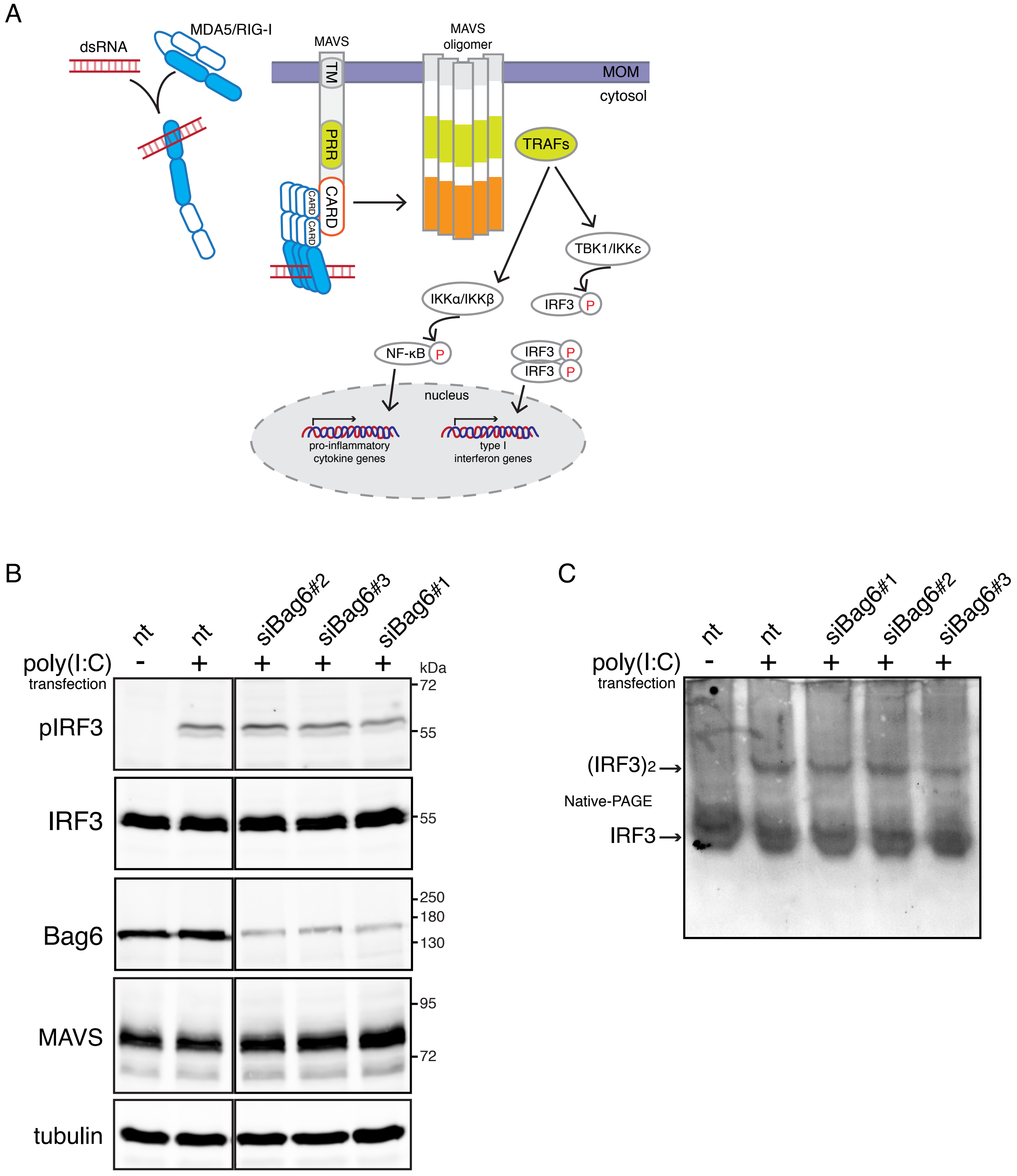
A
dsRNA
MDA5/RIG-I
MAVS
TM
PRR
CARD
CARD
CARD
MAVS
oligomer
MOM
cytosol
TRAFs
TBK1/IKKε
IRF3 P
IKKα/IKKβ
NF-κB P
IRF3 P
IRF3 P
nucleus
pro-inflammatory
cytokine genes
type I
interferon genes
B
nt
nt
siBag6#2
siBag6#3
siBag6#1
poly(I:C)
transfection
- + + + +
kDa
pIRF3
IRF3
Bag6
MAVS
tubulin
72
55
55
250
180
130
95
72
C
nt
nt
siBag6#1
siBag6#2
siBag6#3
poly(I:C)
transfection
- + + + +
(IRF3)2
Native-PAGE
IRF3

**Fig. S6. Bag6 does not modulate poly(I:C)-induced MAVS-dependent signalling.** (A) Schematic of RIG-I-like receptor (RLR)-dependent signalling pathway. Cytosolic double-stranded viral RNA is initially sensed by the host cell's RLRs (RIG-I and MDA5). Upon binding to the viral RNA, RLRs undergo a conformational change that exposes their caspase activation and recruitment domains (CARDs) for interactions with downstream signalling proteins. Modification of CARDs with K63 polyubiquitin chains results in the oligomerisation and activation of RLRs. Activated RLRs are recruited to the mitochondria outer membrane, where they interact with MAVS via their CARDs (Hou et al., 2011). This interaction as well as K63-linked polyubiquitination of MAVS (Liu et al., 2017) induces a conformational change leading to the formation of functional prion-like MAVS aggregates and activation of downstream signalling (Hou et al., 2011). MAVS aggregates form a signalling scaffold by recruiting and activating multiple tumour necrosis factor receptor-associated factors (TRAFs), which in turn recruit kinase complexes that phosphorylate the transcription factors IRF3 and NF-kB leading to the transcriptional upregulation of pro-inflammatory cytokines and type I interferons. (B,C) Bag6 depletion does not affect IRF3 activation in response to cytosolic poly(I:C). Control KO cells transfected with non-targeting (nt) or three independent Bag6-targeting siRNAs for 72 h were mock-treated (-) or treated with poly(I:C) (+) using typical transfection conditions (1 μg/ml, 12 h) (Li et al., 2021; Liu et al., 2017; Zhang et al., 2020) and subsequently analysed by immunoblotting for the indicated endogenous proteins. In (B), all samples were analysed on the same gel and the combined image was produced using the same brightness and contrast settings.

### Figure S7

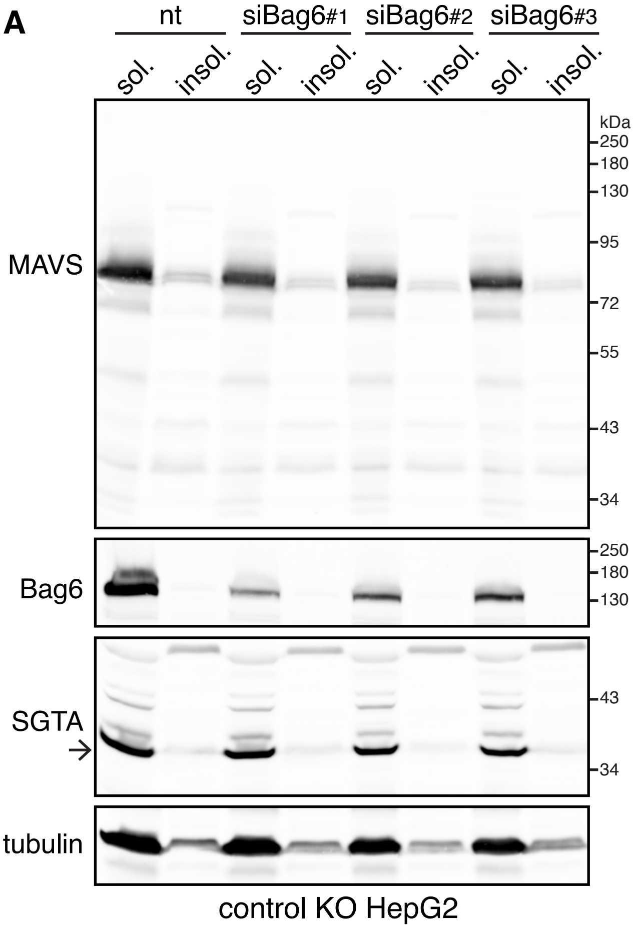

A
nt
siBag6#1
siBag6#2
siBag6#3
sol.
insol.
kDa
250
180
130
95
72
55
43
34
MAVS
Bag6
SGTA
tubulin
control KO HepG2


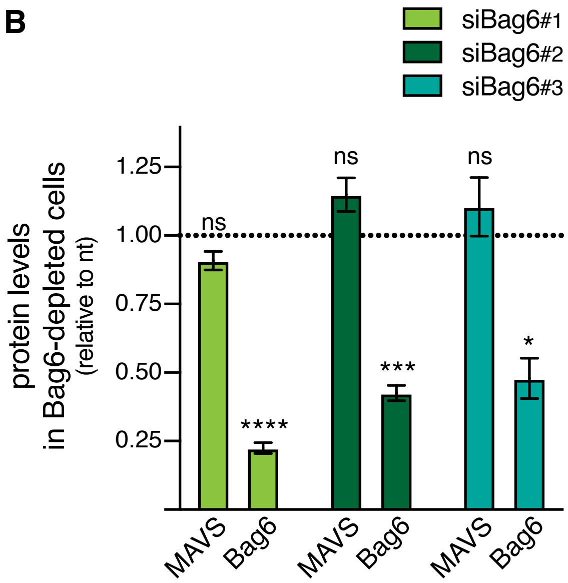

B
siBag6#1
siBag6#2
siBag6#3
protein levels in Bag6-depleted cells
(relative to nt)
1.25
1.00
0.75
0.50
0.25
ns
****
***
*
MAVS
Bag6

**Fig. S7. Bag6 deficiency does not affect steady-state MAVS levels.** (A) Control KO cells treated with non-targeting (nt) or three independent Bag6-targeting siRNAs for 72 h were harvested with a non-ionic detergent-containing lysis buffer. The resulting lysates were separated into soluble and insoluble fractions and analysed by immunoblotting for the indicated endogenous proteins. Note that endogenous MAVS appears to be uniformly soluble in Bag6-depleted cells. SGTA and tubulin remain uniformly soluble under all conditions and serve as fractionation and loading controls. (B) Bag6 and MAVS levels in siBag6-treated cells relative to nt siRNA-treated cells, where protein levels were set to 1. Shown are means ± s.e.m. for four biological replicates as shown in (A). *$P < 0.05$; ***$P < 0.001$; ****$P < 0.0001$; ns, not significant (one-way ANOVA with Dunnett's multiple comparison tests).
